# Histo-molecular differentiation of renal cancer subtypes by mass spectrometry imaging and rapid proteome profiling of formalin-fixed paraffin-embedded tumor tissue sections

**DOI:** 10.1101/2020.02.19.956433

**Authors:** Uwe Möginger, Niels Marcussen, Ole N. Jensen

**Author notes:** Corresponding author: Professor Ole N. Jensen, PhD*, Department of Biochemistry & Molecular Biology and VILLUM Center for Bioanalytical Sciences, University of Southern Denmark, DK-5230 Odense M, Denmark. Present address: Global Research Technologies, Novo Nordisk A/S, Novo Nordisk Park, DK-2760 Måløv, Denmark.

## Abstract

Pathology differentiation of renal cancer types is challenging due to tissue similarities or overlapping histological features of various tumor (sub)types. As assessment is often manually conducted outcomes can be prone to human error and therefore require high-level expertise and experience. Mass spectrometry can provide detailed histo-molecular information on tissue and is becoming increasingly popular in clinical settings. Spatially resolving technologies such as mass spectrometry imaging and quantitative microproteomics profiling in combination with machine learning approaches provide promising tools for automated tumor classification of clinical tissue sections.

In this proof of concept study we used MALDI-MS imaging (MSI) and rapid LC-MS/MS-based microproteomics technologies (15 min/sample) to analyze formalin-fixed paraffin embedded (FFPE) tissue sections and classify renal oncocytoma (RO, n=11), clear cell renal cell carcinoma (ccRCC, n=12) and chromophobe renal cell carcinoma (ChRCC, n=5). Both methods were able to distinguish ccRCC, RO and ChRCC in cross-validation experiments. MSI correctly classified 87% of the patients whereas the rapid LC-MS/MS-based microproteomics approach correctly classified 100% of the patients.

This strategy involving MSI and rapid proteome profiling by LC-MS/MS reveals molecular features of tumor sections and enables cancer subtype classification. Mass spectrometry provides a promising complementary approach to current pathological technologies for precise digitized diagnosis of diseases.

## Introduction

Kidney cancer (renal cell carcinoma, RCC) accounts for 2.2% of all diagnosed cancers and is the 13^th^ most common cause of cancer deaths worldwide [1]. Clear cell renal cell carcinoma (ccRCC) constitutes 70% of all kidney cancers [2] and exhibits the highest rate of metastasis among renal carcinomas. Two other common but less aggressive subtypes of renal carcinoma are chromophobe renal cell carcinoma (ChRCC) and the essentially benign renal oncocytoma (RO), which account for 5% and 3-7 % of all cases, respectively [3, 4]. The ability to distinguish between the malignant cancer types ccRCC and ChRCC and the benign RO is crucial for a patient in terms of prognosis, progression and intervention strategies as severe as total nephrectomy. Histopathological kidney cancer diagnostics faces many challenges in daily routine. Typically, test panels consisting of a combination of different chemical and immuno-histochemical staining methods are used to systematically obtain a diagnosis [5]. Overlapping histological features can make it difficult to differentiate tumor types. Analysis, interpretation and diagnosis/prognosis greatly rely on visual inspection and the experience of the involved clinical pathologists. Complementary techniques such as magnet resonance imaging (MRI) and electron microscopy involve costly instrumentation. Moreover, specific antibodies for staining can be expensive or unavailable for certain molecular targets. Mass spectrometry is emerging as a promising new tool in translational research, from molecular imaging of tissue sections to deep protein profiling of tissue samples [6]. The digital data readout provided by high mass accuracy mass spectrometry and feasibility of molecular quantification makes it a very attractive technology in translational research for investigating human diseases and for diagnostics and prognostics purposes in the clinic. Improvements in mass spectrometry instrument performance and computational analysis paved the way for applications in clinical microbiology [7] and clinical genetics analysis [8]. The fact that mass spectrometry can be applied to a variety of different bio-molecules such as peptides, lipids, nucleic acid makes it extremely versatile and expands the translational and diagnostic possibilities greatly [8–11].

Molecular imaging of tissue sections by MALDI mass spectrometry (MSI) was introduced more than 20 years ago [12, 13] and it has been applied in translational research and clinical applications, to study injuries, diseases, or distinguish between different cancer types such as Pancreatic Ductal Adenocarcinoma or Epithelial Ovarian Cancer Histotypes [14–18].

Mass spectrometry-based proteomics relies on advanced LC-ESI-MS/MS technology, where peptide mixtures are separated by liquid chromatography (LC) prior analysis by electrospray ionization tandem mass spectrometry (ESI MS/MS) and protein identification by protein database searching [19, 20]. Current LC-MS/MS strategies enable comprehensive quantitative protein profiling from tissues and body fluids [21, 22]. While having been used to identify potential biomarkers or new candidate cancer targets and molecular signaling networks the relatively long LC gradients (hours) and extensive sample preparation protocols make it difficult to apply in a routine clinical setting. Modern mass spectrometers are steadily increasing in sensitivity and scanning speed [23]. In addition, improved chromatographic systems that enable rapid solid phase extraction integrated with reproducible separations are emerging [24–27], enabling fast (minutes) and sensitive (nanogram) analysis of complex biological samples.

We hypothesized that histo-molecular information from both MALDI MS imaging (MSI), *in situ* protein digestion and LC-MS/MS applied to detailed characterization of 5 μm cancer formalin fixed paraffin embedded (FFPE) tissue sections will provide spatial molecular maps and sufficiently deep proteome profiles to characterize and classify tumor subtypes. We investigated this by testing a series of malignant and benign renal carcinomas, including clear cell renal cell carcinoma (ccRCC), chromophobe renal cell carcinoma (ChRCC) and renal oncocytoma (RO). We obtained histo-molecular images at a resolution of 150μm × 150μm that sufficed to spatially resolve features to distinguish tumor subtype areas from surrounding tissue. Miniaturized sample preparation by *in situ* protein digestion was used to recover peptides from distinct areas of the FFPE tumor sections for rapid proteome profiling by LC-MS/MS.

## Material and Methods

### Materials

Xylene (analytical grade), ammonium bicarbonate, Sodium citrate, trifluor-acetic acid (TFA), formic acid (FA), acetic acid (AcOH), acetonitrile (ACN), methanol and α-Cyano-4-hydroxycinnamic acid (CHCA) were purchased from Sigma. Polyimide coated fused silica capillary (75 μm ID) was from PostNova, C_18_ Reprosil Pur reversed phase material was from Dr. Maisch (Ammerbuch-Entringe, Germany), recombinant Trypsin was purchased from Promega (WI. USA), Indium-tin-oxide (ITO) glass slides were purchased from Bruker (Bremen, Germany), water was Milli-Q filtered.

### Formalin fixed paraffin embedded samples

Patient samples were collected at Odense University Hospital, Denmark. All samples were obtained upon patient’s consent. Formalin fixed paraffin embedded (FFPE) tissues from 11 RO patients, 12 ccRCC patients and 5 ChRCC patients were used for LC-MSMS analysis (for ChRCC due to the lower number of patients 2 subsequent slides were used from 2 patients adding up to a total of 7 sections). Out of the patient cohort 9 RO, 9 ccRCC and 5 ChRCC were used for mass spectrometry imaging analysis.

### Tissue preparation

#### Preparation of formalin fixed paraffin embedded samples

FFPE blocks were cut into 5 μm thick sections and mounted onto indium tin oxide (ITO) covered glass slides (for MSI) or regular microscopy glass slides (for LC-MS/MS). Before deparaffination slides were left on a heated block at 65° C for 1 hour to improve adhesion (an overview on the used FFPE samples can be found in supplementary table S1 and S2).

#### Deparaffination

FFPE section slides were incubated in Xylene for an initial 10 min. and then another 5 min. using fresh solution each time. Slides were shortly dipped into 96% EtOH before they were washed for 2 min in a mixture of chloroform/Ethanol/AcOH (3:6:1; v:v:v). The slides were then washed in 96% EtOH, 70% EtOH, 50% EtOH and Water for 30 sec. each.

#### Antigen retrieval

Tissue slides were heated in 10mM citric acid buffer pH 6 for 10 min in a microwave oven at 400 Watt (just below the boiling point) before left for further 60 min incubation at 98°C on a heating plate. Slides were cooled down to room temperature and incubated for 5 minutes in 25 mM ammonium bicarbonate (ABC) buffer. Slides were allowed to dry before application of trypsin protease.

#### Tryptic digest

##### For MALDI MS imaging

20μg of Trypsin (Promega) was used per slide and was dissolved at a concentration of 100ng/μl in 25mM ABC /10% ACN before being deposited on the tissue using the iMatrixSpray [28] device equipped with a heating bed (Tardo Gmbh, Subingen, Switzerland) using the following settings: sprayer height = 70mm, speed = 70mm/s, density = 1μL/cm^2^, line distance= 1 mm, gas pressure= 2.5 bar, heat bed temperature= 25°C . After trypsin deposition the slides were incubated in a humid chamber containing 10mM ABC/ 50% MeOH at 37°C over night.

##### For *on-tissue* digest intended for LC-MS/MS proteome profiling

Droplets of 2μl Trypsin solution (50ng/μL in 25mM ABC /10%ACN, 0.02%SDS) were deposited using a gel loading pipet tip. Droplets were placed on 2-6 different tumor areas of each FFPE tissue section. The extraction positions were chosen randomly within the defined tumor margins which were defined by HE-stain and MSI clustering. The droplets were quickly allowed to dry to prevent spreading across the tissue. Slides were transferred to a closed humidity chamber (10mM ABC /50% MeOH) for overnight digestion at 37°C. The digestion spots were extracted twice with 2μL of 0.1% FA and twice with 1.5μL of 30%ACN. Fractions were combined for each sample and speedvac dried. Samples were reconstituted in 0.05%TFA and shortly spun down prior injection into the LC-MS system.

#### Matrix application

Matrix solutions were freshly prepared from recrystallized α-cyano-4-hydroxycinnamic acid (CHCA) matrix (10mg/mL in 50% Acetonitrile 1% TFA). Matrix was sprayed using the iMatrixSpray (Tardo, Switzerland). Temperature of the heatbed was set at 25°C. The sprayer distance was set to 70mm. Spray speed was set to 100 mm/s. Matrix was sprayed in 3 rounds: 8 cycles with a flowrate of 0.5μl/cm^2^ line distance of 1mm, 8 cycles of 1μl/cm^2^ line distance of 1mm, 8 cycles of 1μl/cm^2^ and a line distance of 2mm.

#### MALDI MS Imaging data acquisition

Optical images of the tissue were obtained before matrix application using a flatbed scanner (Epson) at resolutions of 2400dpi. The imaging data was acquired via FlexImaging software (Bruker, Daltonics, Bremen, version 3.1) with 500 shots/ pixel on a Ultraflextreme MALDI-TOF/TOF MS (Bruker Daltonics, Bremen) equipped with a SmartBeam laser (Nd:YAG 355 nm). External mass calibration was performed with a tryptic digest of bovine serum albumin (Sigma). Spatial resolution was set to 150μm in x- and y-direction. Mass spectra were acquired in positive ion reflector mode in the range *m/z* 600-3500. (An average sum spectrum of each cancer condition can be found in Supplementary material 1: Figure S1)

#### LC-MS/MS analysis

LC-MS/MS data was acquired by an Orbitrap Q-Exactive HF-X (Thermo, Bremen) coupled to an Ultimate 3000 capillary flow LC-system. Setup was modified from Thermo Scientific Technical note: 72827. Peptide samples were loaded at 150μl/min (2% ACN, 0.05% TFA) for 30 sec onto a 5μm, 0.3 × 5 mm, Acclaim PepMap trapping cartridge (Thermo Scientific). Samples were then eluted onto a pulled emitter analytical column (75μm ID, 15cm). The analytical column was “flash-packed” [29] with C_18_ Reprosil Pur resin (3μm) and connected by Nanoviper fittings and a reducing metal union (Valco, Houston, TX). The flowrate of the 15 min gradient was 1.2 μL/min with solvent A: 0.1% formic acid (FA) and solvent B: 0.1% FA in 80% ACN. Gradient conditions for solvent B were as followed: 8% to 25% in 10 min, 25% to 45% in 1.7 min. The trapping cartridge and the analytical column were washed for 1 min at 99%B before returning to initial conditions. The column was equilibrated for 2 min. MS settings: ESI spray voltage 2kV, cap temp=275°C, Resolution: 60k, micro scans =1, max IT =100 ms, AGC =3×10^6^, MSMS resolution 15k, n= top 5, max IT =100 ms, AGC = 1×10^5^.

#### Data Processing of MALDI MS imaging data

The data was baseline subtracted, TIC normalized and statistically recalibrated and then exported into imzML format [30] using the export function of FlexImaging software (Bruker). The exported mass range was m/z 600-3000 with a binning size of 9600 data points. The imzML files were imported into the R environment (version: 3.4.1) and further processed and analyzed using the R MSI package: Cardinal (version: 2.0.3 & 2.4) [31]. In order to extract pixels of tumor tissue each sample was preprocessed as follows: peaklist was generated by peak picking in every 10^th^ spectrum and subsequent peak alignment. The whole data was then resampled using the “height” option and the previous created peaklist as spectrum reference. PCA scores were plotted using car-package (version 3.0.6). Samples were clustered using spatial shrunken centroid clustering [32]. Subsequently, clusters were compared to tumor regions in HE-stained tissue sections (supplementary material 1: Figure S2). The respective clusters containing tumor areas were extracted, so that result files predominantly contained data from tumor areas. The obtained coordinates were then used to extract the corresponding pixel from the unprocessed imzML file. Each tumor type was assigned with a diagnosis factor (ccRCC, RO or ChRCC), which was later used as y-argument in the cross-validation. All extracted imaging acquisition files were further restricted to a mass range of m/z 700-2500. Data was resampled with step size 0.25 Da to allow combining them into one file for further processing. Classification and cross-validation were performed using partial least square discriminant analysis (PLS-DA) [33]. PLS components were tested for optimum with 34 components (**supplementary material 1: Figure S3**). Classification diagnosis was based on the highest scoring condition. Differences between 2 conditions had to be higher than 10% of the highest score to be considered distinguishable)

#### LC-MS/MS data processing

The MaxQuant [34] software package (version 1.5.7.0) was used for protein identification and label-free protein quantitation. LC-MS/MS data was searched against the Swissprot human proteome database, using standard settings and “match between runs” enabled.

Data filtering, processing and statistical analysis of the MaxQuant output files was performed using the Perseus [35] framework (version 1.6.1.3). Data was filtered excluding the following hits: only identified by site, contaminants and reversed. The log-transformed data was filtered for proteins present in at least 70% of all experiments. Significance filtering was based on ANOVA testing, using FDR threshold of 0.01 with Benjamini Hochberg correction. In order to perform PCA analysis and classification missing values were imputed by normal distribution (separately for each column/sample). Data shown in heatmap was Z-score normalized. Perseus output tables were transferred into ClustVis [36] for visualization of hierarchical clustering and principle compound analysis (PCA). Gene Ontology and functional analysis was performed via String DB (version 11.0.0) [37] and Panther DB (version 14.1) [38]. For Panther DB analysis background genome was the human genome and the total of identified proteins from all LC-MSMS runs in the experiments (supplementary material 4-8). Feature optimization cross-validation type was “n-fold” with n = 5. Kernel was either linear or RGF. All other settings were left on their default value.

## Results

In this study we investigated the utility of mass spectrometry-based methods for histo-molecular profiling applications in clinical renal cancer pathology. We analyzed thin tissue/tumor sections from three different renal cancer types (ccRCC, RO, ChRCC) by MALDI MS imaging and by an optimized rapid LC-MS/MS workflow adjusted to suit the demands for clinical settings.

### Imaging by MALDI mass spectrometry

All samples were prepared as 5 μm thin FFPE tissue/tumor sections. The entire FFPE tissue section was analyzed by imaging MALDI MS imaging (MSI). The data was subsequently processed by unsupervised clustering (spatial shrunken centroid clustering [32]). The clustering results **(Figure 1A and 1B)** illustrate the heterogeneity of the tissue sections coming from various tissue types such as stroma, fibrotic, fatty or healthy tissue and the capabilities of imaging MSI for the delineation of cancerous and non-cancerous tissue. Furthermore, when comparing the tumor area of the HE-stain/microscopy with the results from the mass spectrometry imaging based clustering, spectral differences even within the tumor tissue itself can be observed (**Figure 1A and 1C**).

**Figure 1.**
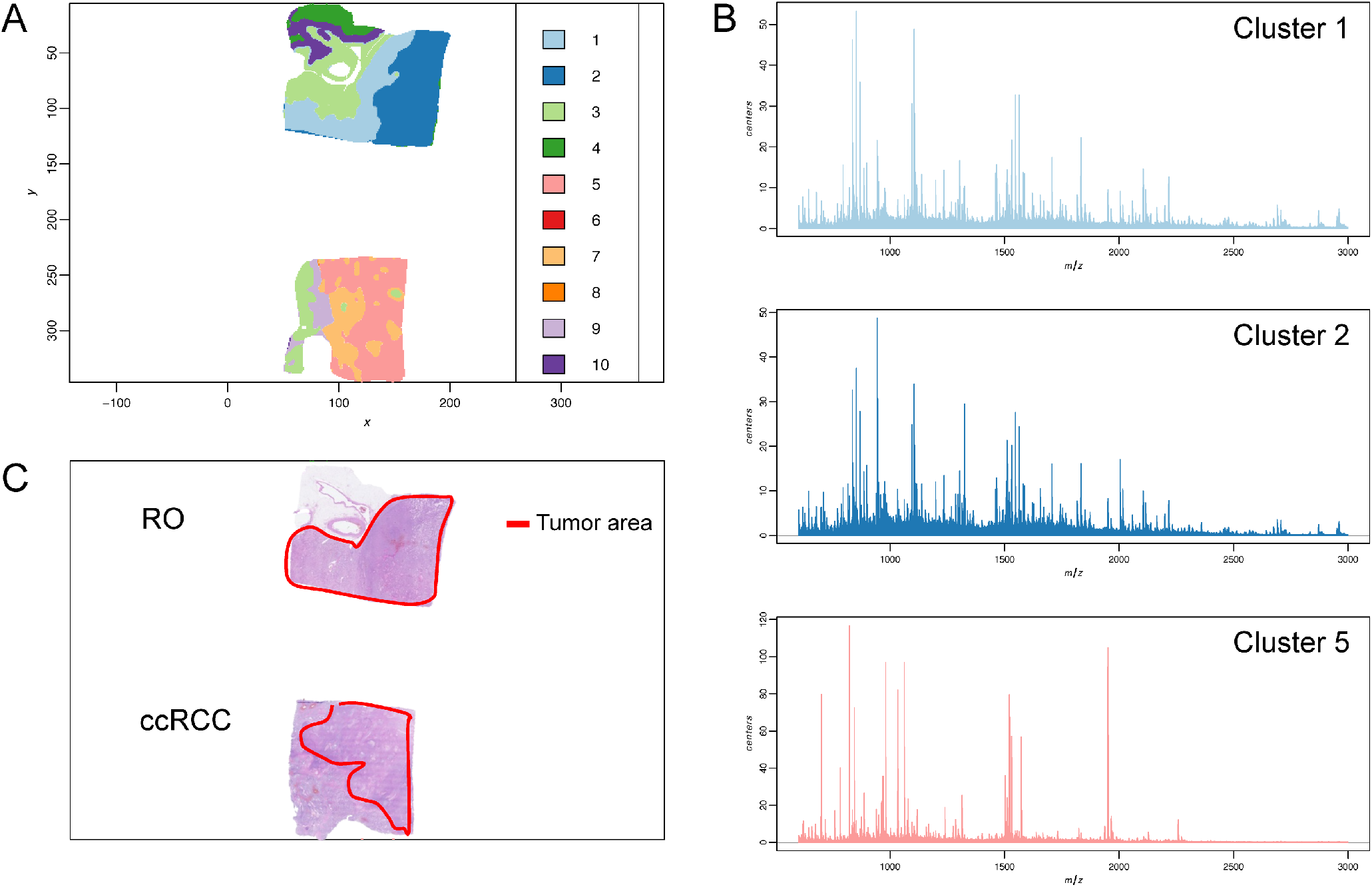
Tumor sample heterogeneity is revealed by mass spectrometry imaging and unsupervised clustering. A) Spatial Shrunken centroid clustering of ccRCC and RO data obtained by imaging mass spectrometry of ccRCC and RO tissue sections. Based on differences and similarities in the spectra each pixel was automatically assigned a certain cluster (indicated by a different color). B) Average MALDI mass spectra of the respective tumor areas (histo-molecular clusters) reveal distinct features and individual variations in the m/z signals. C) HE-stain of tumor tissue section from same FFPE block. Tumor area is indicated in red.

Guided by the unsupervised clustering outcome and the corresponding image obtained by HE-staining, pixels from non-relevant surrounding tissue were discarded and only pixel clusters containing actual tumor tissue were used for subsequent comparative analyses (schematic workflow overview can be found in supplementary material 1 : Figure S4).

In mass spectrometry imaging, principal component analysis (PCA) is often used for initial analysis of a given data set. Variance and similarities within the image sample set were estimated by PCA over the first 3 components. From a pathology viewpoint RO and ChRCC are more difficult to distinguish than ccRCC and ChRCC. As the sample holder for the imaging experiments can only hold 2 slides at a time, we first compared two conditions in a pairwise manner: 9 ccRCC vs. 9 RO **(Figure 2A)** and 5 RO vs. 5 ChRCC **(Figure 2B)**. Then the data set was combined to compare all three cancer conditions (9 ccRCC, 9 RO, 5 ChRCC) to each other **(Figure 2C).**PCA using the first three principle components separate ccRCC well from RO and ChRCC (**Figure 2A, 2C).** Data points from ccRCC showed a wide spread and were splitting into 2 sub-populations. In contrast, the data from RO and ChRCC samples cluster in a much tighter manner and with some overlap **(Figure 2A, 2B, 2C)**. This is particularly the case when considering all three cancer types together (**Figure 2C)**. When compared in a pairwise manner RO and ChRCC show slight separation **(Figure 2B)** suggesting at least some degree of histo-molecular differences between these cancer types. Some overlapping data points in the different cancer type datasets can be observed indicating histo-molecular spectral similarity in parts of the patient tumor tissues. The spread of ccRCC data points in PCA, as compared to the RO and ChRCC subtypes, suggests a greater heterogeneity among the ccRCC patient samples (also observed by LC-MS/MS, see later section).

**Figure 2:**
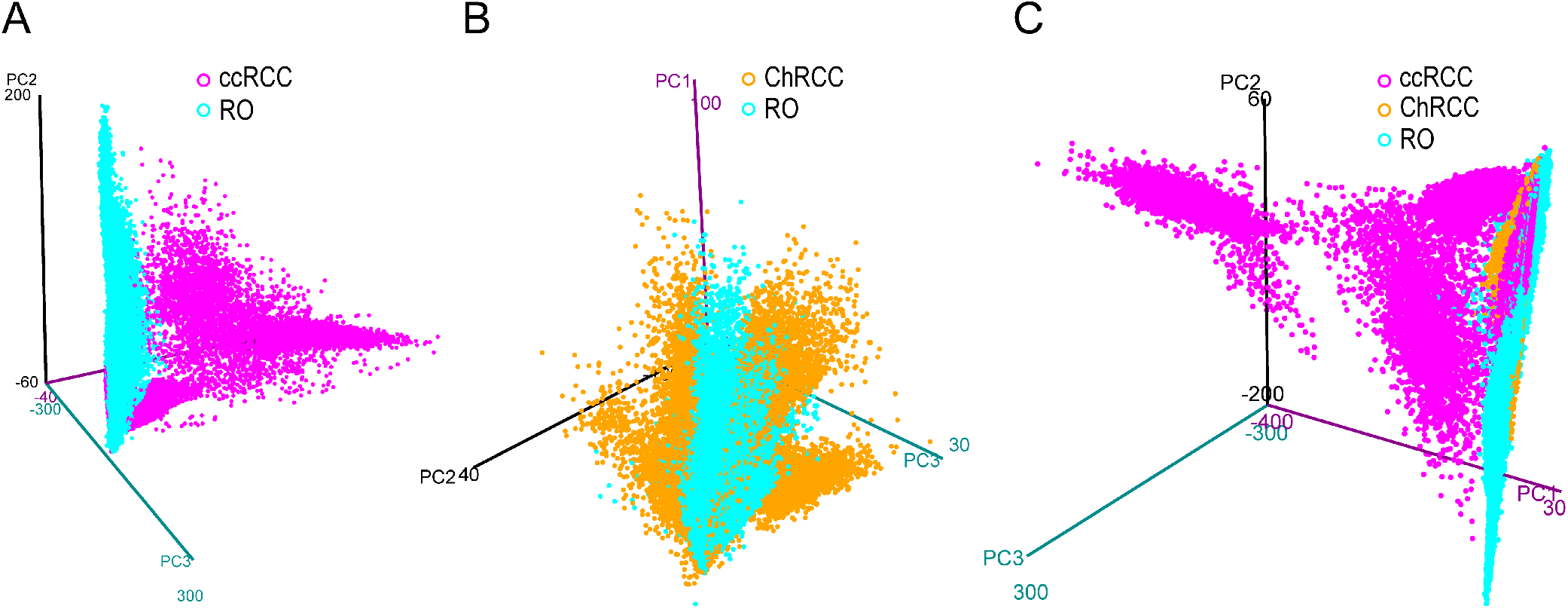
3D PCA score plot from imaging MALDI MS experiments of kidney tumor tissues. Each plot contains the extracted pixel data from all patients of a given cancer type. Data from ccRCC (A) (magenta) and ChRCC (B) (blue) are compared to RO (yellow). (C) Data from all three cancer types are compared to each other. The graph displays the first 3 principle components (PC1, PC2, PC3) plotted against each other. Clear separation of data points between ccRCC and RO can be observed by pairwise comparison but also in the combined comparison to both RO and ChRCC. In pairwise comparison (B) RO and ChRCC show slight separation but exhibit a great number of overlapping features. ccRCC exhibits the largest differences to RO and ChRCC. RO and ChRCC appear to share more spectral similarities.

Next, we assessed the ability of the MSI data to distinguish and classify renal cancer subtypes. We generated a classifier based on partial least squares discriminant analysis (PLS-DA) that can then be applied to a given MSI sample set. Due to the limited number of FFPE kidney tumor samples we chose to use a cross-validation strategy that maximizes the use of a sample set for model generation and testing. In this approach a classifier is trained with imaging data from all samples, except for one sample that is set aside. As this sample is not part of the classifier model it can then be used for testing purposes. This was repeated as many times as there are samples ultimately allowing for testing the complete dataset (for n samples we obtain n classifiers and n tested samples).

The optimized PLS-DA model resulted in an accuracy of cancer subtype prediction of 93% for ccRCC and 88% for RO and for ChRCC (pixel based value). Results of the cross-validation study using PLS-DA to classify 23 kidney tumor samples are depicted in **Figure 3**. The PLS-DA prediction scores for each of the three possible tumor type outcomes are shown, i.e. ccRCC, RO and ChRCC. (Median values and boxplot representation of scores are provided in **supplementary material 1: Figure S5 and table S3**). The scores obtained for each pixel are presented by intensity scaled colors plotted over the respective x-y-coordinate of the tissue/tumor sections.

**Figure 3:**
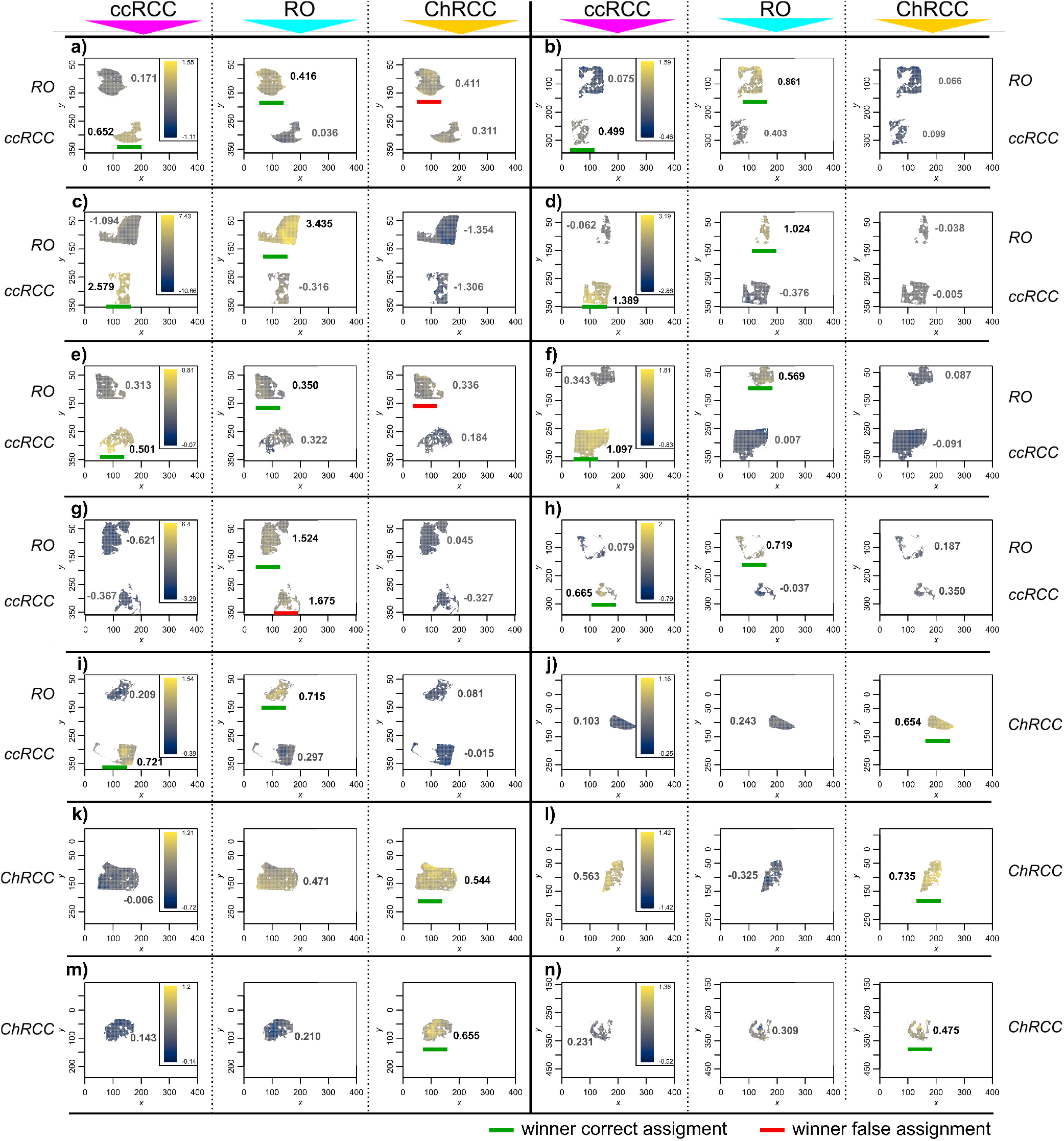
Tumor classification by MALDI MS imaging and cross-validation using PLS-DA classification. Unit of x- and y-axis equals step size (150μm). Classification was performed on extracted pixels/spectra from tumor areas only. Classification of 9 ccRCC and 9 RO (a-i) sample as well as 5 RO sample and 5 ChRCC sample (j-n). Pathology diagnosis of the respective patient samples are indicated to the left and right of the images (*RO, ccRCC* or *ChRCC*). Each spectrum-containing pixel is predicted individually. The prediction scores are represented by a color scale. Each patient sample was scored for the 3 cancer conditions resulting in 3 panels for each condition. Each of the panels displays scores for ccRCC (first panel) RO (second panel) and ChRCC (third panel). The respective testing condition is indicated on top above the panels. Median score for the respective condition is indicated in each panel next to the tissue. Classification is based on the condition achieving the highest score within the 3 predictions. Differences below 10% of the respective highest score were considered too close to be distinguishable. A overview table with median values as well as a boxplot representation of scores is provided in supplementary material 1 Figure S5 and table S3. (Cardinal’s smooth.image-function was used for better visibility. Unprocessed image can be found in supplementary material: 1 Figure S7) Each sample is predominantly predicted in the correct diagnosis, achieving accuracies (pixel-based value) of 93% (ccRCC), 88% (RO), 88% (ChRCC). Winner of the classification is marked with a green bar for correct classification and a red bar for incorrect classification.

Twenty of the 23 patient tumor samples (assignment based on median value) were correctly assigned by the PLS-DA model showing highest intensity and median score for the respective cancer condition **(Figure 3)**. Eight out of nine ccRCC samples were correctly assigned **(Figure 3a-i).** The PLS-DA classification provided high scores for ccRCC samples and clearly distinguished the ccRCC samples from the other two kidney tumor types (Figure 3, left panels). This is in accordance with the PCA results. Likewise, the PLS-DA model provided low scores for ccRCC in the cases of RO and ChRCC samples (Figure 3, middle and right panel). One ccRCC sample was incorrectly classified by PLS-DA as RO **(Figure 3g**).

All 5 ChRCC samples (**Figure 3 j-n**) were correctly assigned having the highest score for the ChRCC condition (right panel). PCA indicated mass spectral similarities between the RO and ChRCC samples. Likewise, the PLS-DA model reflects such similarities in the classification outcome. Two ChRCC patient samples received highest scores for ChRCC but only slightly lower scores for RO (**Figure 3k, 3n**). Furthermore, in the case of two RO sample **(Figure 3a, 3e)** the classification could exclude ccRCC as diagnosis. Although the highest median score was correctly achieved for RO (Supplementary material 1: **table S3**) the difference between RO and ChRCC was considered as too small (<10% of respective max. median score) for a clear distinction of these two tumor types. Notably one kidney tumor sample (Figure 3l) exhibited a unusual scoring pattern as compared to the other tumor samples. This particular sample received high scores for both ccRCC and ChRCC classification (ChRCC being the highest). As mentioned above, we typically observed clear distinction between ccRCC and ChRCC in all the other cases. Upon further pathology and microproteomics analysis this tumor section was re-classified as a sarcomatoid transformation (see below), i.e. a tumor type not included in the PLS-DA model used for classification.

The relative importance of individual histo-molecular features of the classifier can be visualized by plotting the PLS coefficients for each condition as a function of m/z values (**supplementary material 1: Figure S6**). A positive coefficient indicates presence or higher abundance of the m/z value in the respective cancer model. A negative coefficient indicates absence or lower abundance in the respective condition. For ccRCC the two highest-ranking m/z values were m/z=723.5 and m/z=704.5. The two highest values for RO were m/z=806.5 and m/z=1640.0 whereas the most influential signals for ChRCC comprised m/z= 1169.5 and m/z=1039.5 (top 100 list of the features can be found in **supplementary material 2).** Unfortunately, we were not able to obtain informative MALDI MS/MS fragment ion spectra in order to reveal the identity of these peptide ion signals. Nevertheless, for classification purposes the knowledge of distinct protein/peptide identities (m/z values) is not necessary as long as the signal is characteristic for the tested condition.

In conclusion mass spectrometry imaging provided histo-molecular tumor profiles that can be used to distinguish renal cancer subtypes. However, the misclassification of one ccRCC patient and uncertainty of two additional diagnosis outcomes suggested that additional independent test methods would be beneficial for confident classification of renal cancer tumor types.

### LC-MS/MS based rapid proteome profiling of tumor sections

MALDI MS imaging provides spatial resolution that is helpful to address molecular heterogeneity in tissue sections. However, MALDI MS imaging lacks analytical depth due to the limited dynamic range of MALDI TOF MS and the poor performance of MALDI MS/MS for protein identification by peptide sequencing directly from tissue sections. Deeper insight into the tissue and tumor histo-molecular profiles and their variance will provide more diagnostic features. We therefore adapted and optimized a microproteomics approach, combining *in situ* protein sample preparation with fast label-free proteome profiling LC-MS/MS. First a miniaturized *in situ* sample preparation method was applied where a small droplet of trypsin solution is placed directly onto the tumor area of interest within a thin tissue section. After overnight incubation the digested protein extract from the tumor area is subsequently recovered and analyzed by mass spectrometry [39]. We reduced the LC-MS/MS analysis time from 90 minutes to 15 min by using short LC gradients and rapid MS/MS functions, allowing for a sample throughput of up to 80 samples per day. A total of 125 *in situ* extracted areas from renal tumor sections were analyzed. Two to six *in situ* extracts were taken from each renal tumor sample (11 RO sections from 11 patients: 47 extraction spots; 12 ccRCC sections from 12 patients: 49 extraction spots; 7 ChRCC sections from 5 patients: 29 extraction spots). Fast label-free LC-MS/MS based microproteomics analysis of all 125 *in situ* digested tumor areas resulted in a total of 2124 identified human proteins. We filtered the data for proteins that were present in at least 70 % of all samples thereby reducing the protein number to 412 proteins. Comparative data analysis was performed for proteins that were significantly altered (FDR=0.01) in any of the renal cancer subtypes resulting in a list of 346 differentially regulated proteins. We then used unsupervised hierarchical clustering and PCA to identify similarities and differences between the tumor samples. The x-axis dendrogram of the heatmap shows that the majority of the renal tumor samples grouped according to cancer subtype RO, ccRCC or ChRCC **(Figure 4A)**. Several large clusters of “co-regulated” proteins are evident on the y-axis dendrogram and heatmap for the individual cancer subtypes. This clearly demonstrates that there are renal cancer subtype specific histo-molecular features and patterns in the microproteomics dataset.

**Figure 4:**
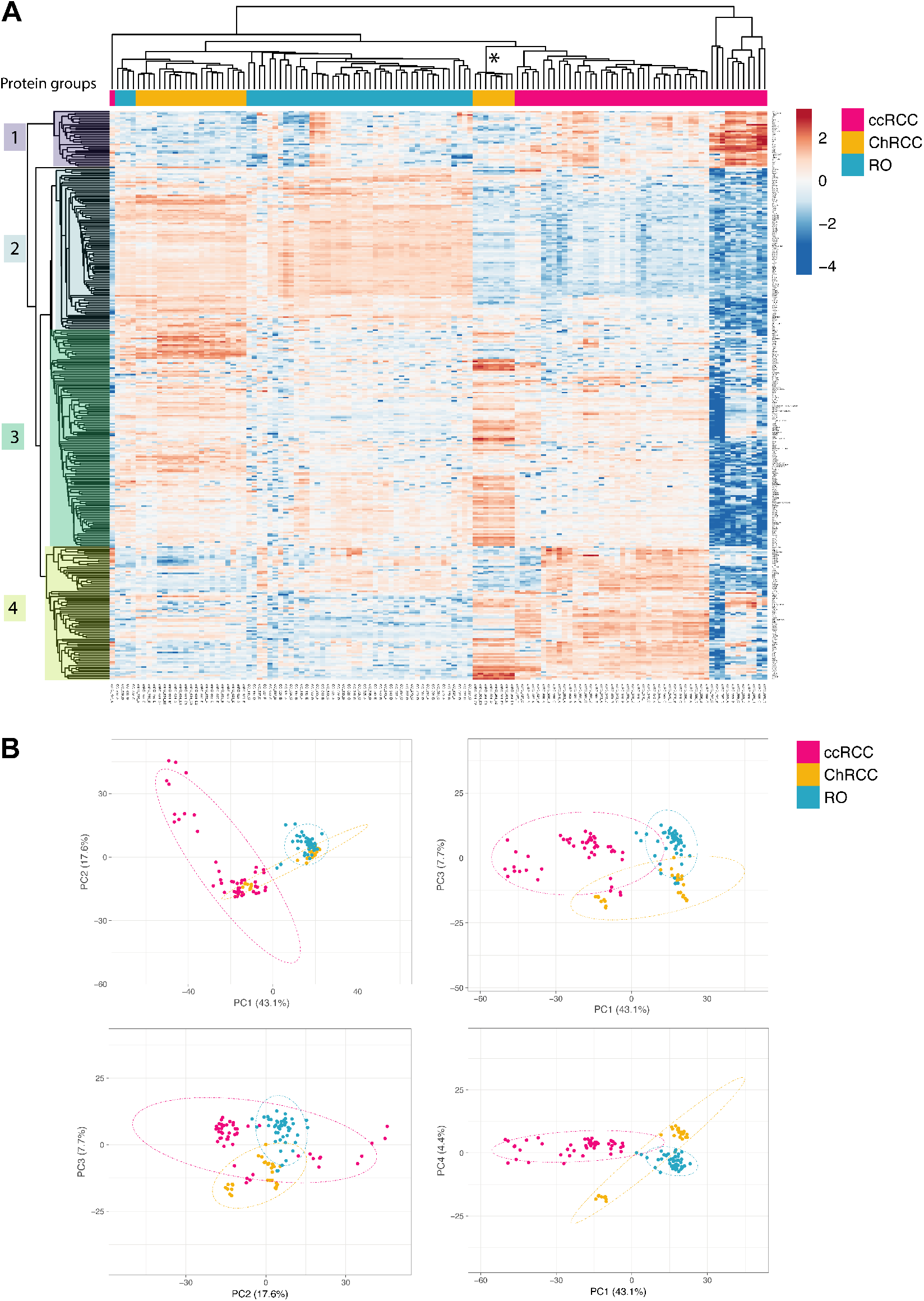
Unsupervised renal cancer subtype classification by microproteomics using rapid LC-MS/MS protein profiling. A) Heatmap and hierarchical clustering of differential relative protein abundances. Columns indicate samples and rows indicate proteins. The renal cancer subtype of the patient sample is indicated in colored bars on top. The graph shows the large similarities in protein expression profiles among patient samples with the same cancer subtype causing them to cluster together. Furthermore, hierarchical clustering of the protein abundances reveals protein cluster that are detected in a cancer subtype specific manner. Protein groups selected for subsequent network analysis are indicated by color blocks on the y-axis dendrogram (groups 1-4).The asterisk * marks outlier patient from sample figure 3l. B) Principal component analysis of the sample set. Dotted ellipses are such that with a probability of 95% a new observation from the same group will fall inside the area. The first (PC1) and second (PC2) component explain 17.6 % of the total variance whereas the other components lie at 7.7% and 4.4% respectively. There is a clear separation of ccRCC and RO samples already in the first two principal components. Differences between RO and ChRCC are subtle and are only evident when considering components that display lower variance (PC2:PC3 and PC3:PC4). The small group of the eight ChRCC-derived sarcomatoid renal cancers samples cluster relatively far from the other ChRCC samples, thereby identifying these as clear “outliers” that require further attention.

The protein expression profiles of the three renal cancer subtypes are different based on the heatmap patterns. ccRCC clearly differs from RO and ChRCC (**Figure 4A**: Protein group 2 and 4). RO and ChRCC display some differences but generally exhibit a more similar expression pattern (**Figure 4A**: Protein Group 2).

These differences and similarities were also revealed by PCA analysis of the microproteomics dataset. RO and ChRCC separate clearly from ccRCC **(Figure 4B)**. RO and ChRCC datapoints are located close together, indicating that differences between the RO and ChRCC cancer subtypes are less dominant. When considering principal components exhibiting less variance (PC3 and PC4), separation of RO and ChRCC sample data is observed (**Figure 4B**).

We observed eight ChRCC proteomics datasets that separated clearly form the other ChRCC datasets, both in hierarchical clustering analysis (**Figure 4A**) and PCA (**Figure 4B**). The protein expression profile of these 8 samples exhibited some similarities to both ChRCC and ccRCC. Interestingly, this data originated from a tumor from a single patient. This was the same patient that also exhibited outlier MSI data with similarities to both ChRCC and ccRCC tumor types, as discussed above (Figure 3l). Further pathology analysis revealed that these samples were sarcomatoid renal cancer, originating from ChRCC and, thus, indeed different from the other ChRCC samples.

### Protein differences in cancer subtypes

Hierarchical clustering of the proteomics datasets revealed major differences in relative protein abundance between the three renal cancer tumor types. (**Figure 4A**). We investigated the nature of these histo-molecular differences by examining the correlation of these proteins to cellular structures, functions, or biochemical processes. Protein groups that exhibited distinctive abundances for the respective cancer type (**Figure 4:** ccRCC: group 1 & 4, RO: group 2, ChRCC: group 3) were searched for their involvement in protein interaction networks (supplementary material 9) as well as for their functional roles by using gene ontology (GO) enrichment (**Figure 5**, supplementary material 3-8). We compared GO enrichment relative to the experimental gene background as well as to the complete human genome (Figure 5, human genome background: red). The experimental gene background contained all genes corresponding to all 2124 identified proteins in the LC-MSMS experiments (Figure 5, experimental background: blue).

**Figure 5:**
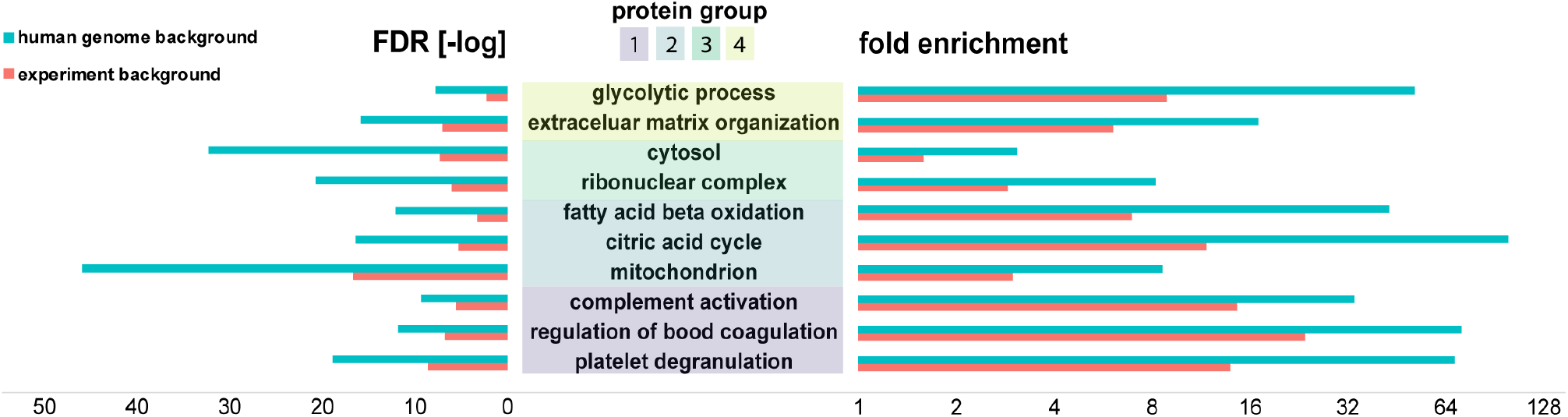
Bioinformatics analysis (PantherDB) identified enriched biochemical functions in renal tumors. Protein groups were compared against a background of all the 2124 proteins identified in the experiment (blue) and against the background of the human genome (red). Fold enrichment (increase over expected value) as well as −log of the false discovery rate (FDR) are shown.

RO and ChRCC exhibited a set of upregulated proteins (**Figure 4A**, *protein group 2)* that were enriched for mitochondria associated proteins (GO:0005739), including various ATP synthase subunits. Enriched protein functions comprised *oxidative phosphorylation* (hsa00190), *citrate cycle* (hsa00020), and *fatty acid beta oxidation* (GO:0006635).

ChRCC-specific regulated proteins (**Figure 4A**, *protein group 3*) included cytoplasmic proteins (GO:0044444), and proteins associated with *cytoplasmic vesicles* (GO:0031982) and *ribonucleoprotein complexes* (GO:1990904).

Subtype-specific protein groups in ccRCC (**Figure 4A**, *protein group 1, 4*) were functionally enriched for *complement activation* (GO:0006956), *regulation of blood coagulation* (GO:0030193) and *platelet degranulation* (GO:0002576). Functions of *protein group 4* were linked with *extra cellular matrix organization* (GO:0043062) and *cytoskeletal binding* (GO:0008092) including proteins *collagen* and *laminin*. We also found several proteins such as *glyceraldehyde-3-phosphate dehydrogenase* associated with the *glycolytic process* (GO:0006096).

These functionally important findings can be correlated to known biochemical and morphological features of each of the renal cancer subtypes. It is known that the number of mitochondria is increased in RO and ChRCC tumors (e.g. increased oxidative phosphorylation) [40]. It is also known that ccRCC contains a highly vascularized stroma (complement, coagulation, etc.) and exhibits a strong Warburg effect (glycolysis) [41]. Large intracellular vesicles are found in ChRCC (cytoplasmic proteins, vesicle proteins) [40].

### Classification

Unsupervised data analysis demonstrated the presence of renal cancer subtype specific differences in the tumor protein profiles. Next, we investigated the feasibility of tumor classification by using the microproteomics data to train a prediction algorithm. We implemented the tumor classification model by using a support vector machine (SVM) approach. The sarcomatoid sample was excluded from the classification. We chose the k-fold cross validation strategy [42] (“n-fold” in Perseus). Here the data is randomly distributed in k groups. The model was then trained with data from k-1 groups and the prediction was applied to the samples in the remaining group. This was repeated k times. Low k-values tend to overestimate error rates. In our study 2-6 extraction spots (samples) were derived from an FFPE section from each patient so too high k-values could underestimate the true error rate. We therefore tested the prediction error rate over several k-values (**Figure 6 A**) applying Radial Basis Function (RBF) and linear kernel functions [43]. As imputation could have an effect on the classification outcome we compared performance to a classification without imputation using the proteins that were present in all sample (100% valid values=27 proteins). For 70% valid values the tested error rates were in the range 0-3.4% for linear (4 wrong predictions at k=2, linear kernel) and 0-2.6% (3 wrong predictions at k=2, RBF kernel). However, k=2 is a very low k-value (excluding half of the samples from the training set) and the error rate is most likely overestimated in this case. For more commonly used k-values (k=3-10) the error rate was 1.7% (2 incorrect predictions) at the highest for k=3 and 0% for any other k-value. Incorrectly predicted outcomes included samples from one RO patient that was predicted as ccRCC. Classification with the data set using 100% valid values without imputation showed error rates of 0-6% (linear kernel) and 0-3.5% (RBF kernel). Over higher k-values the error rate was 0-3.4% and 0-0.8%, respectively. Error rates were slightly higher than for 70% valid values. Given the low protein number used for the classification the outcome was surprisingly positive. In both cases 70% and 100% valid values we observed RBF performing overall slightly better than linear kernel. As error rates for both valid values were quite comparable, we concluded that in our case the imputation did not heavily bias the outcome.

**Figure 6:**
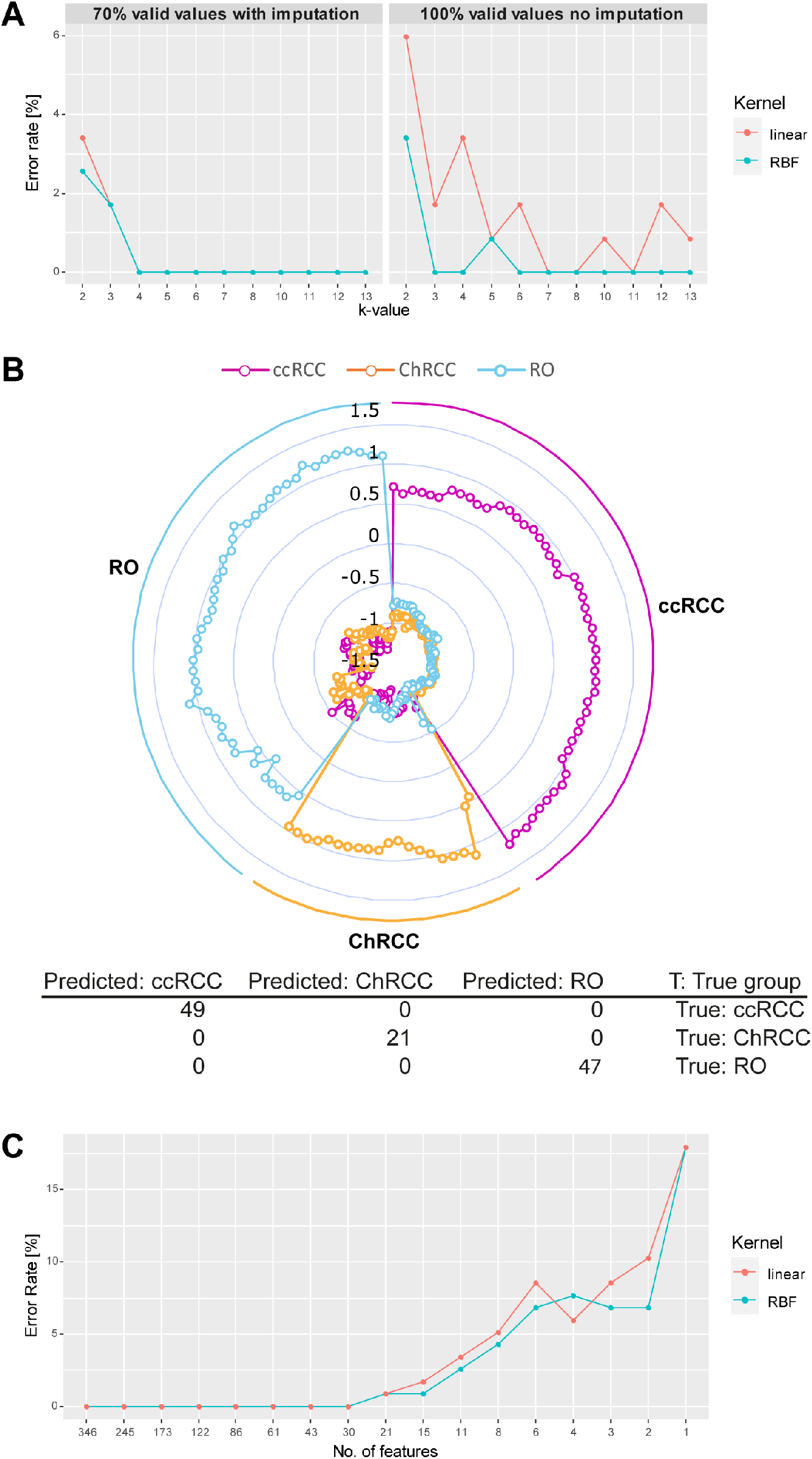
Microproteomics and SVM model correctly classifies all renal tumor subtypes. A) Development of cross validation classification error rate in relation to increasing k-value (division of sample set in k groups. k-1 groups will be used for training and 1 group for testing). Results of 70%valid values using imputation of missing values (right) as well as 100%valid values without imputation (left) are shown. Two classes of algorithms were compared: RBF and linear kernel. RBF performs slightly better than the linear kernel function with lower error rates. Values chosen for k>=3, error rates vary between 0% and 1.6% (2 wrong prediction out of 117) for both tested kernel. B) Radar plot of the cross-validated classification (k=5, kernel=RBF) of proteome profiles obtained from each extraction spot sample. Each sample is plotted equi-angular around the center. The pathological diagnosis (ground truth) for each sample at its angular position is indicated on the outside of the radar plot (all ccRCC samples: right, all ChRCC samples:bottom, all RO samples:right). A given sample is represented by 3 datapoints (dots) plotted on a straight axis originating from the center. Each of the 3 datapoints represents the classification score for one of the 3 cancer types (scores for ccRCC: magenta, ChRCC: yellow, RO: blue). Scores range from lowest (center) to highest (outer circle).The highest score indicates highest likelihood for the respective cancertype. The plot shows that for all samples the cancer type with the highest score correlates with the respective pathological diagnosis, indicating the high accuracy of the classification. C) Feature optimization. The error rate for linear kernel and RBF are plotted over the number of ranked features (proteins). Decreasing feature number results in increase of false predictions. Minimum number of features for 0% error rate is at 30 for both RBF and linear kernel (list of ranked proteins can be found in supplementary material 10).

Figure 6B exemplifies the outcome of the cross-validation resulted for RBF and k=5 (around 23 samples per group equivalent to 4-5 patients excluded from the training set). Each sample was scored for the three tested conditions (ccRCC, RO, ChRCC). The highest scoring condition was used to classify a given sample. Results are shown in a radar plot (**Figure 6B**) and demonstrate 100% accuracy in prediction of renal cancer subtypes.

We initially used all 346 differentially abundant histo-molecular features (proteins) to classify the tumor subtypes. Next, we sought to estimate the minimum number of features that suffice to correctly classify all the renal tumor samples (for k=5 and RBF). We used the feature optimization function in the Perseus software, which first ranks the features and then tests the error rate for a decreasing number of features (**Figure 6C**). The minimal number out of the 346 features was found to be 30 features (list of the ranked proteins can be found in supplementary material 9). Further reducing the number to 21 features resulted in an error rate of 0.8% and as little as 4 features lead to an error rate of 7.7%. Conclusively only a portion of the dataset, would suffice to successfully classify all the kidney tumor samples, which reflects also in the low error rates of the 100% valid values (**Figure 6A)** using only 27 proteins (**supplementary material 3**). However, keeping an excess of quantified protein features would be beneficial as “safety margin” assuring a high enough number of quantified protein features for robust classification of tumors.

### Data integration from MSI and rapid proteome profiling

Having both MSI and microproteomics sets of data at hand provides several advantages for classification of cancer tumor FFPE samples. Using the MSI approach for tumor classification we observed a higher error rate than with the rapid micro-proteomics approach. In two cases MSI could exclude one cancer type but was not providing clear results towards an RO or a ChRCC diagnosis. Another case where one ccRCC sample was misclassified as RO is particularly problematic as ccRCC might need surgery whereas RO does not. Therefore, by integrating the micro-proteomics classification data the outcome of the MSI classification can be further confirmed, clarified or rejected (**Table 1**). This allows more confidence in diagnosis or could possibly even provide further information on cancer stage or treatment strategies. In a case where the classification model does not cover the cancer condition such as in the patient sample with sarcomatoid transformation we have demonstrated how irregularities and inconsistencies are detected by both MSI and rapid LC-MS/MS based microproteomics (**Figure 3l, 5A, 5B**). This provides an opportunity to further investigate, refine and expand the range of computational and statistical classification models.

**Table 1:**
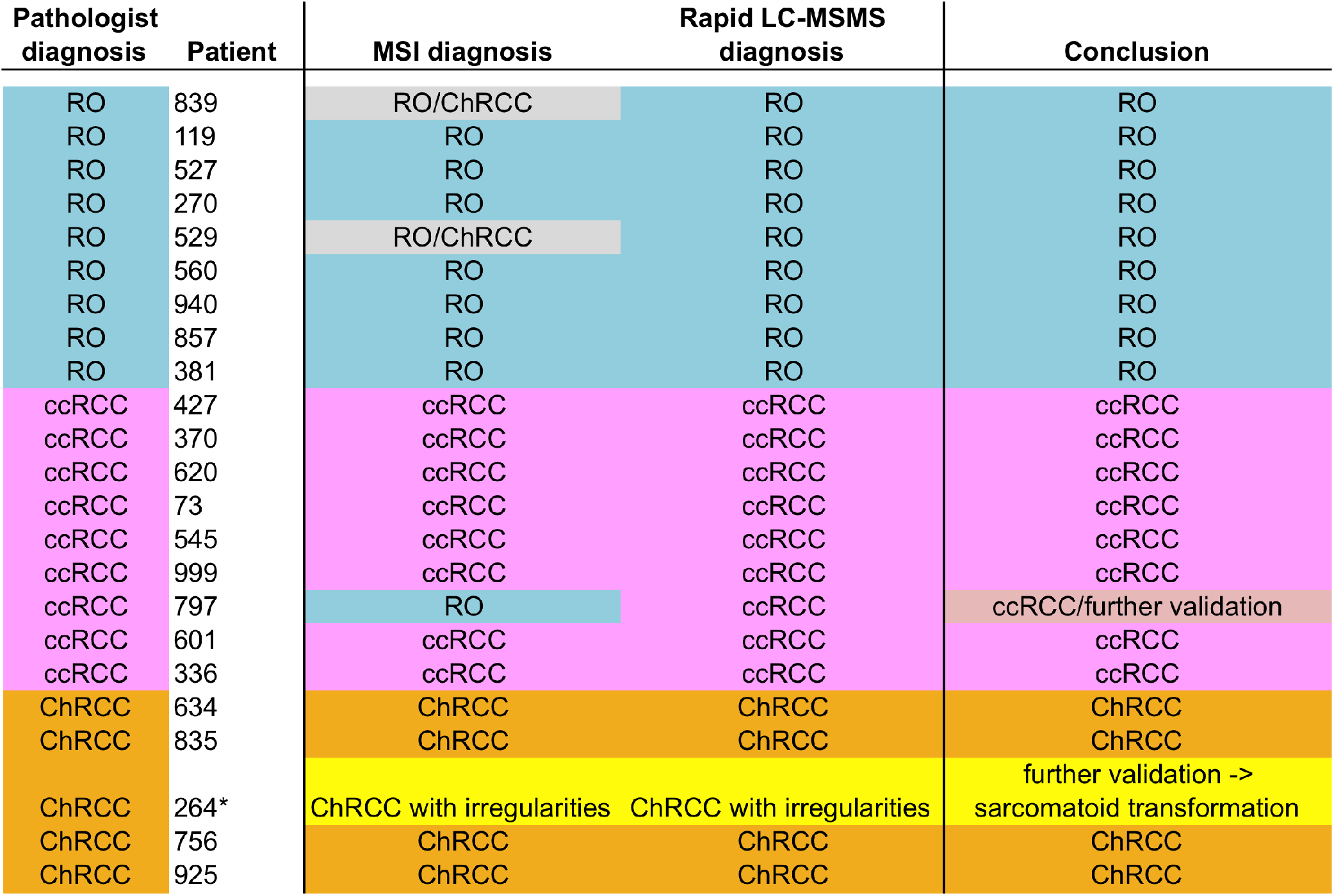
integrated testing strategy for classification of renal cancer types. Initial pathologist diagnosis and patient number are indicated in the first 2 columns. *Patient sample showed irregularities and after reassessment could be diagnosed as sarcomatoid transformation. Concluding contradictory results would either necessitate further validation or the outcome of the more reliable method (LC-MSMS) could be favored

An additional application making use of the combined data set includes the investigation of histo-molecular properties observed in MSI (e.g. intra- tumor heterogeneity) by correlation to information from the rapid microproteomics approach. Usually a detailed investigation of MSI feature data is achieved by either microproteomics “in situ protein digestion” or laser microdissection based approaches [44] using LC-MS/MS based proteomics analysis with long LC gradient times (1-4 hours). Despite the shorter gradient times and thus lower protein coverage in the present work the information can nevertheless be used to investigate histo-molecular properties observed in MSI (e.g. intra-tumor heterogeneity). We exemplified this in figure 7, using the RO MSI data set previously shown (**Figure 1,** top). Unsupervised spatial shrunken centroid clustering of MSI data [32] revealed two distinct regions within the tumor area (**Figure 7A**: cluster 1 and cluster 2). Correlating the LC-MSMS data from the respective extraction spots within these distinct regions in deed reveals significant differential abundances in 80 proteins (**Figure 7B**, the list of proteins can be found in **supplementary material 11**). Hierarchical clustering (**Figure 7C**) of these 80 proteins with regard to their extraction position correlates well with the distinct regions depicted by the MSI clustering (MSI Cluster 2 correlates with extraction spots e-f, MSI Cluster 2 correlates with extraction spots a-d; **Figure 7A**). The proteomics data suggests a lower abundance of mitochondrial associated proteins and a higher abundance in some cytoskeletal protein binding proteins in cluster 2 (**Figure 7D**). The area comprising Cluster 2 located on the edge of the tumor and might indicate the differences that can be encountered between the inner and outer tumor regions [45].

**Figure 7:**
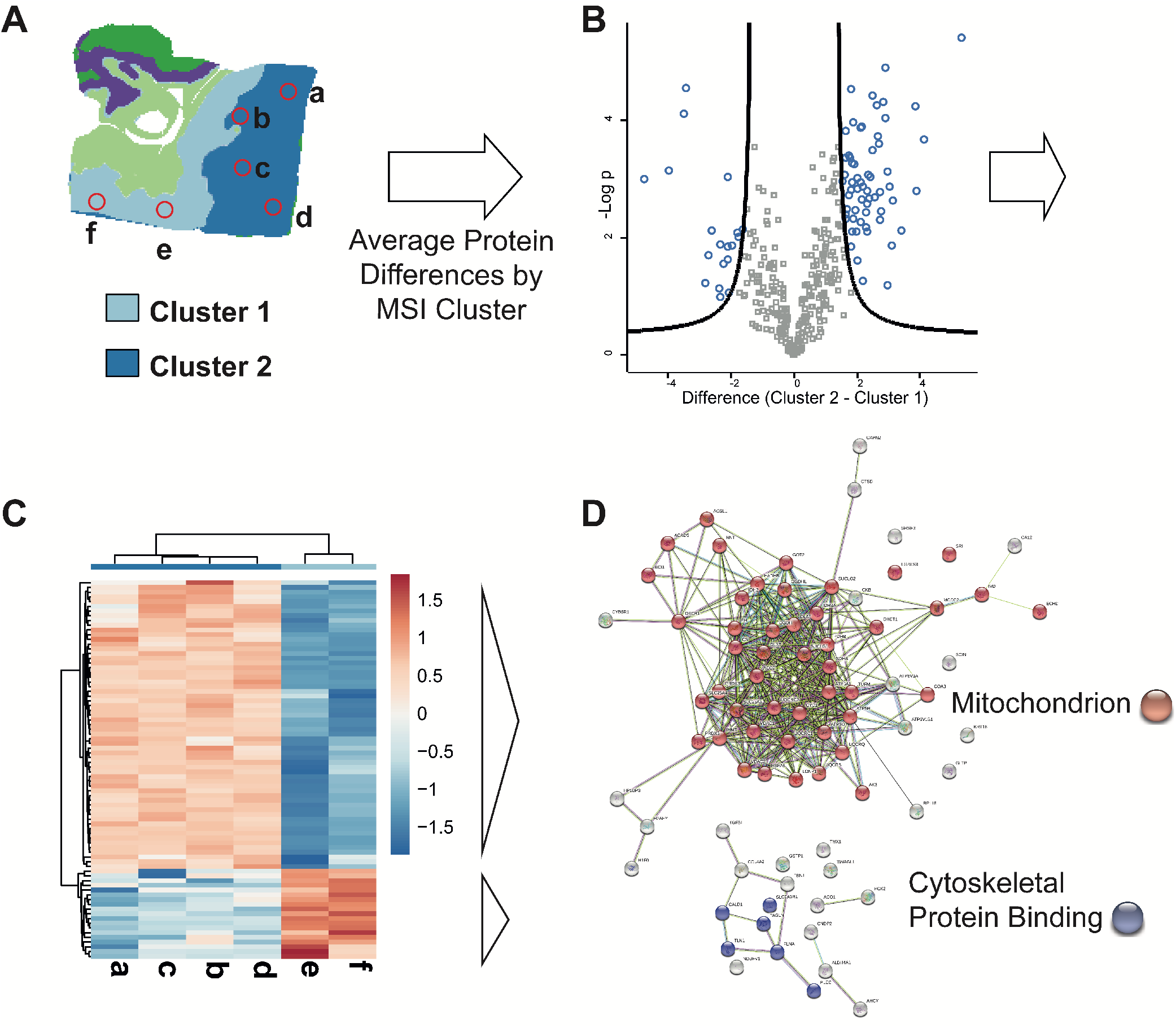
Combined use of MS imaging and rapid LC-MSMS microproteomics provides histo-molecular details of tumor heterogeneity. A) MSI based unsupervised clustering analysis of a RO patient sample. Two clusters (cluster 1 and cluster 2) are detected within the tumor area. Positions used for extraction of LC-MSMS samples are indicated by red circles extractions a-f (2 extractions for cluster 1, 4 extractions in cluster 2). B) Volcano plot of LC-MSMS data derived ratio of protein abundance (Cluster 2 / Cluster1). The –log p values of protein abundances are plotted over the difference of the protein abundance. The black line depicts the chosen significance threshold (p=0.01, 2-fold difference). Proteins above the thresholds are colored in blue and mark significant differences in protein abundance among the 2 compared regions. Proteins with increased abundance are found on the right side of the plot, proteins with decreased abundance are found on the left side of the plot. C) Heatmap display of the significantly different proteins from extraction spots a-f. On the x-axis extraction spots a-d and e-f group together by hierarchical clustering. The grouping is in correlation to the MSI clustering data. On the y axis two protein groups can be observed distinguishing the two x-axis-cluster. One group is upregulated in Cluster 1 the other group is upregulated in Cluster 2. D) StringDB network analysis of upregulated proteins in MSI-Cluster-2 (top, sample: a-d) and upregulated proteins in MSI-Cluster 1 (bottom, sample: e-f). For a given set of proteins the string data base (www.string-db.org) provides information on a dataset in terms of protein attributes such as interaction, function or cellular occurrence. Proteins with higher abundance in Cluster 2 are mainly mitochondrial associated proteins whereas cluster 1 shows increase abundance of cytoskeletal protein binding proteins.

## Discussion

The increasing incidence of renal cancer in western countries calls for improved technologies for detection, diagnosis, treatment and prognosis. Innovative mass spectrometry-based applications are beginning to address challenges in clinics and the healthcare sector, such as the use of targeted proteomics to characterize noninvasive liquid biopsies [46] or the so called iKnife, enabling surgeons to identify cancerous tissue in real time [47, 48]. Mass spectrometry is becoming increasingly applicable in a clinical setting [49, 50]. FFPE sections are a valuable source for mass spectrometry-based diagnosis. As many of the sample preparation steps for MS analysis overlap with the preparation steps for (immuno)histochemical staining, they can be seamlessly fit into the high-throughput sample preparation pipeline for FFPE sections (deparaffination, antigen retrieval) already existing in many hospitals.

Our proof of concept study demonstrates the potential and benefits of mass spectrometry techniques for detailed characterization of clinical specimen. Specifically, we demonstrate that mass spectrometry provides valuable results in the diagnosis of different renal cancer subtypes (ccRCC, RO and ChRCC). The imaging mass spectrometry (MSI) approach allows to collect spatially resolved spectra without *a priori* knowledge of the tissue, thereby enabling the differentiation between cancerous and noncancerous tissue, as well as subtyping of tumors.

Earlier large scale MSI classification studies have demonstrated results with accuracies ranging from 81% to nearly 100% in subtyping non-small cell lung cancer [51], classifying primary lung and pancreatic cancer [52] as well as differentiating between 6 common cancer types (esophagus, breast, colon, liver, stomach, thyroid gland) [53]. In our study MALDI-MSI could diagnose 87% (20 out of 23) of the tested patients correctly. It has to be pointed out that when transferring our study into a larger scale (n>100 samples) misclassification rates are expected to increase. In two of the 3 misclassified cases it was possible to narrow down the diagnosis to either RO or ChRCC. Despite the promising results the misclassification of one ccRCC sample as RO might be problematic since RO may not require surgery but ccRCC does. Both cases stress how using rapid proteome profiling data in parallel provided additional confidence and can help avoid a false negative prognosis.

Both MSI and LC-MS/MS PCA data showed that the patient-to-patient tumor variability is significant for ccRCC. Possible reasons might be due to necrotic areas or increased bleeding observed in some of the tissues. Furthermore we did not consider difference in grades, which might have an influence on the data spread. For robust MSI performance inclusion of a larger patient cohort (n>100) will likely provide higher confidence and resolve this issue or even provide differentiation of tumor grades.

LC-MS/MS based microproteomics analysis correctly classified all tested renal tumor samples in cross validation experiments. The efficient peptide separation and sequencing capability of LC-MS/MS provided deeper insight into the renal cancer proteome than possible by the MSI approach alone. Remarkably, unsupervised clustering identified data inconsistencies and irregularities in the patient cohort. An unexpected feature pattern revealed a sarcomatoid transformation within the ChRCC cohort, without *a priori* knowledge (**Figure 4A, 4B**). This goes to demonstrate that once the “digital” data is acquired then the computational and statistical applications can uncover relevant and important features of the patient datasets. This sensitivity, specificity and versatility will have major implications for future clinical practices, including histo-molecular pathology technologies.

Using short LC runs of only 15 min. we generated a list of 346 significantly altered proteins (p=0.01). The minimum number of proteins determined to be necessary for 100% accurate tumor classification was much lower (30 features). This low number of features enables a targeted proteomics approach aimed at quantifying only a select panel of proteins. Using fewer features would also allow further reduction of LC run time and increase overall sample throughput. Using our fast LC-MS/MS setup we analyzed a total of 125 samples in a series without experiencing blocking of the LC columns, glass capillaries or ESI needles. LC systems such as the EvoSep system [25] that are specifically dedicated for clinical applications and tailored to be used also by non LC-MS experts can add additional robustness to this approach. Furthermore, implementation of image pattern guided pipetting robots may enhance reproducibility and throughput, e.g. using liquid extraction surface analysis (LESA) technology [54, 55]. The latter has been successfully applied in the study of traumatic brain injuries [56] as well as in mouse brain for the identification of proteins and peptides from MSI experiments [57]. The missing value problem is still a common problem in label free quantitative proteomics. Successful implementation of protein identification on MS1 level only has been presented recently [58] and could be interesting in the here presented context to follow up in future experiments.

Functional protein analysis using bioinformatics tools revealed molecular networks and biochemical processes consistent with previously known macroscopic, morphological and histological features of the renal cancer subtypes. RO and ChRCC exhibited upregulation of mitochondrial associated proteins. Increased numbers of mitochondria are frequently observed in these cancer types by electron microscopy [59] and have been identified in previous proteomics studies [60]. As most cancer rely on glycolysis (Warburg effect) this seems rather unusual. However, those mitochondria are dysfunctional and it has been speculated that the increase in number might be a cellular compensation response [61].

In addition increased intracytoplasmic associated proteins were detected in ChRCC distinguishing it from the other cancer types. Microscopically, ChRCC distinguishes from other renal carcinomas by its pale cytoplasm resulting from large intracytoplasmic vesicles explaining the relative increase of intracellular cytoplasm-associated proteins and vesicle proteins.

Clear cell renal cell carcinoma frequently contains zones of hemorrhage that are most likely responsible for the increased levels of complement and coagulation cascade associated proteins, as determined by our microproteomics method. ccRCC is also characterized by hypervascular stroma [3], which may account for the enrichment of extracellular matrix proteins. Enhanced glycolysis as a hallmark of many cancer types including ccRCC [41] correlating well with our detection of upregulated glycolysis associated proteins.

For classification we applied PLS-DA to MSI data and support vector machine to the LC-MSMS data. These common classification methods have previously been applied to MSI for the differentiation of papillary and renal cell carcinoma based on lipidomics analysis [62] as well as for the classification of epithelial ovarian cancer subtypes [16]. There are, however, numerous other classification methods available. Mascini *et al.* used principal component linear discriminant analysis in order to predict treatment response in xenograft models of triple-negative breast cancer [63]. Recently, deep convolutional networks have been proposed [64].

Both MSI and short gradient LC-MS/MS microproteomics methods come with their individual advantages. Applying both approaches in parallel for routine analysis is most beneficial to improve confidence in diagnosis and identify irregularities. In order to create very robust classifiers for use in clinical settings the promising results of this study need to be further supported in the future by analysis of larger patient cohorts.

With the enormous progress in sample handling and instrument technology, machine learning [65] and the availability of new databases [66] mass spectrometry is on its way to become a versatile tool in the hospital clinics of the future.

## Acknowledgements and Funding

Proteomics and mass spectrometry research at SDU are supported by generous grants to the VILLUM Center for Bioanalytical Sciences (VILLUM Foundation grant no. 7292 to O.N.J.) and PRO-MS: Danish National Mass Spectrometry Platform for Functional Proteomics (grant no. 5072-00007B to O.N.J.). We thank Veit Schwämmle and Livia Rosa Fernandes for advice and discussion on the project.

## Authors Contributions

U.M., O.N.J. and N.M. planned and outlined the project. N.M. provided the patient samples and patient diagnosis. U.M. performed all experiments and data analysis. U.M. and O.N.J. wrote the manuscript.

