## Supplementary material 1 for "Histo-molecular differentiation of renal cancer subtypes by mass spectrometry imaging and rapid proteome profiling of formalin-fixed paraffin-embedded tumor tissue sections"

### Supplementary table S1

**Kidney tissue samples**

**FFPE**

| Cancer type | clear cell renal carcinoma no. (ccRCC) | Grade | Gender | Leibovich score | Age | | Renal Oncocytoma no. (RO) | Grade | Gender | Age | | chromophobe renal carcinoma no. (ChRCC) | Grade | Gender | Age |
|---|---|---|---|---|---|---|---|---|---|---|---|---|---|---|---|
| | 427 | 4 | M | 8 | 69 | | 270 | | M | 73 | | 835 | 3 | M | 63 |
| | 797 | 2 | F | 3 | 69 | | 381 | | M | 79 | | 634 | 3 | F | 73 |
| | 370 | 3 | M | 5 | 59 | | 119 | | F | 51 | | 756 | 3 | F | 26 |
| | 073 | 2 | M | 2 | 82 | | 857 | | F | 68 | | 923 | 2 | M | 44 |
| | 999 | 2 | M | 5 | 50 | | 527 | | M | 82 | | 264 | 4 | M | 39 |
| | 545 | 2 | F | 2 | 43 | | 940 | | F | 70 | | | | | |
| | 620 | 4 | M | 9 | 63 | | 839 | | F | 76 | | | | | |
| | 336 | 3 | M | 4 | 75 | | 560 | | M | 66 | | | | | |
| | 601 | 2 | M | 2 | 58 | | 529 | | F | 73 | | | | | |
| | 930 | 2 | M | 3 | 48 | | 924 | | M | 52 | | | | | |
| | 310 | 2 | F | 0 | 58 | | 725 | | F | 55 | | | | | |
| | 853 | 2 | M | 0 | 63 | | | | | | | | | | |

Overview of FFPE sample number used in this study sorted according to the cancer subtype diagnosis (ccRCC, RO, ChRCC).

**Supplementary table S2:**
**MSI sample overview**

<table>
<tr><th>IMS Figure 3</th><th>top</th><th>bottom</th></tr>
<tr><td>a</td><td>839</td><td>427</td></tr>
<tr><td>b</td><td>119</td><td>370</td></tr>
<tr><td>c</td><td>527</td><td>620</td></tr>
<tr><td>d</td><td>270</td><td>073</td></tr>
<tr><td>e</td><td>529</td><td>545</td></tr>
<tr><td>f</td><td>560</td><td>999</td></tr>
<tr><td>g</td><td>940</td><td>797</td></tr>
<tr><td>h</td><td>857</td><td>601</td></tr>
<tr><td>i</td><td>381</td><td>336</td></tr>
<tr><td>j</td><td colspan="2">634</td></tr>
<tr><td>k</td><td colspan="2">835</td></tr>
<tr><td>l</td><td colspan="2">264</td></tr>
<tr><td>m</td><td colspan="2">756</td></tr>
<tr><td>n</td><td colspan="2">923</td></tr>
</table>

Sample overview on patient samples used for MALDI MSI analysis.

#### Supplementary Figure S1: Averaged MALDI spectra of respective cancer types

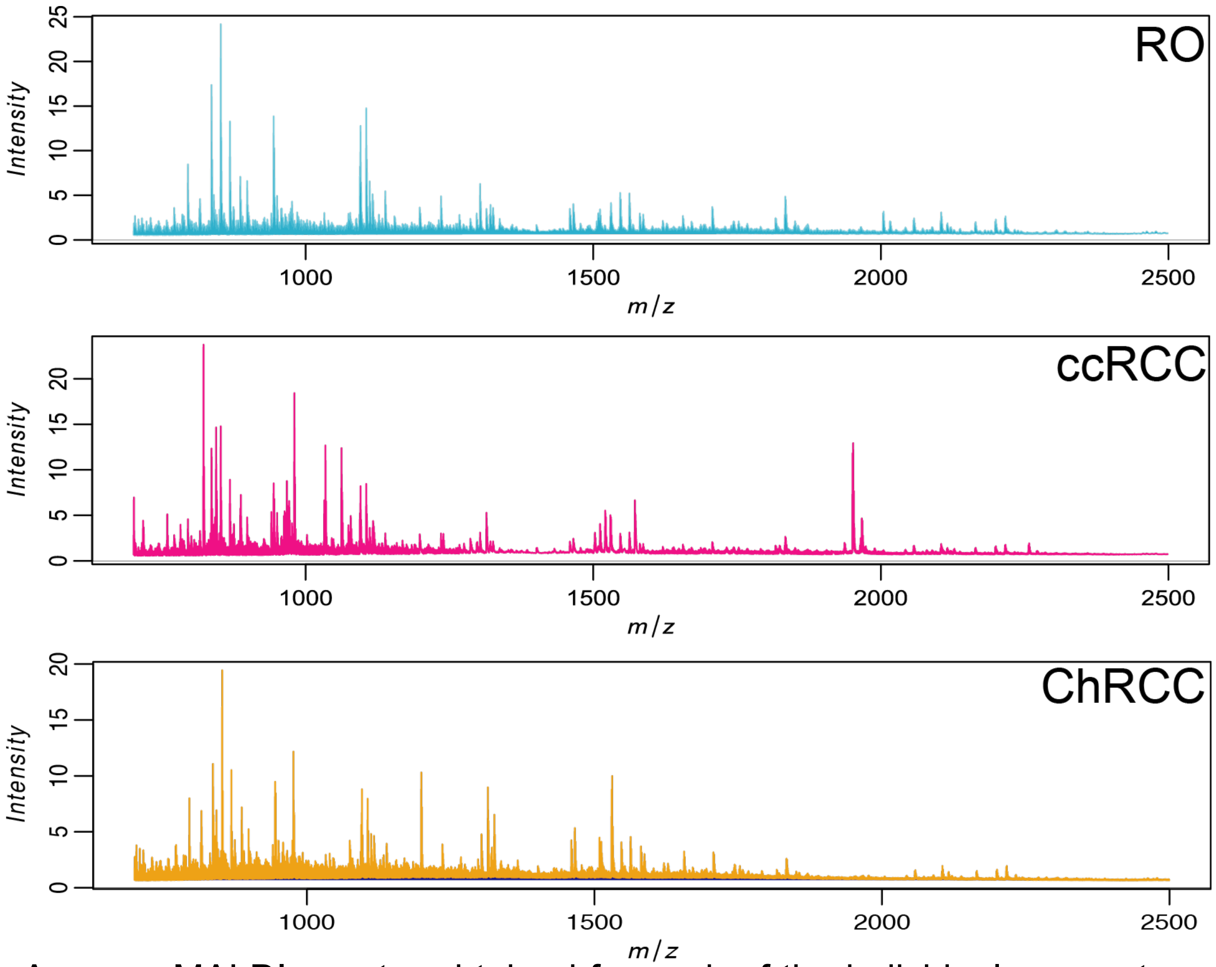


Average MALDI spectra obtained for each of the individual cancer types. Average spectra are based on the extracted pixel. The spectra of the 3 cancer types show differences in their average spectra. however as the pixel amount/patient sample is varying obvious differences in the average spectra can be misleading when looking for distinctive features.

**Supplementary Figure S2:** HE-stains of FFPE sections used in this study

Indicates tumor area

Site of extraction (Note the size of the circle is not correlated to the actual extraction area)

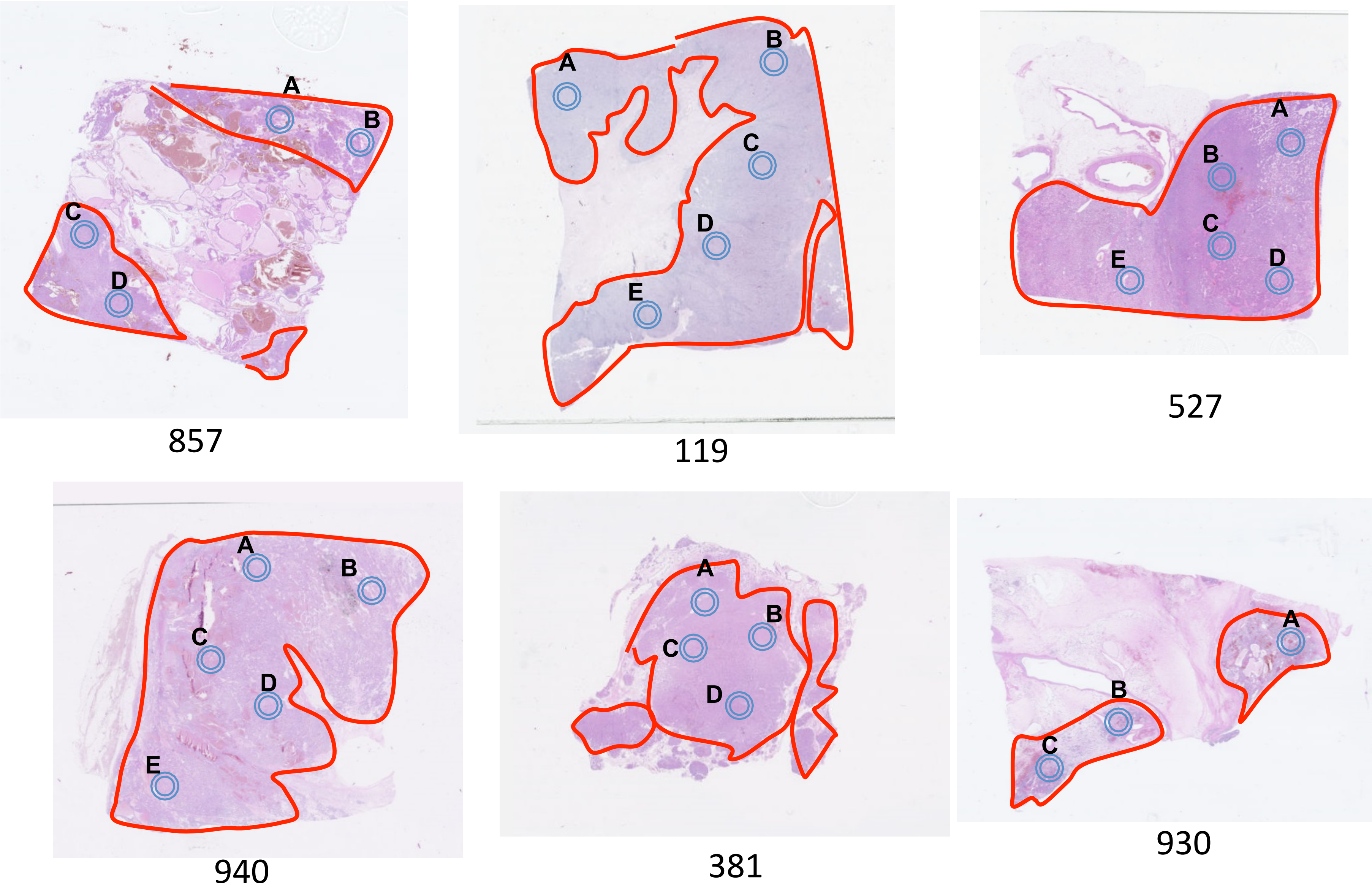

Indicates tumor area

Site of extraction  (Note the size of the circle is not correlated to the actual extraction area)

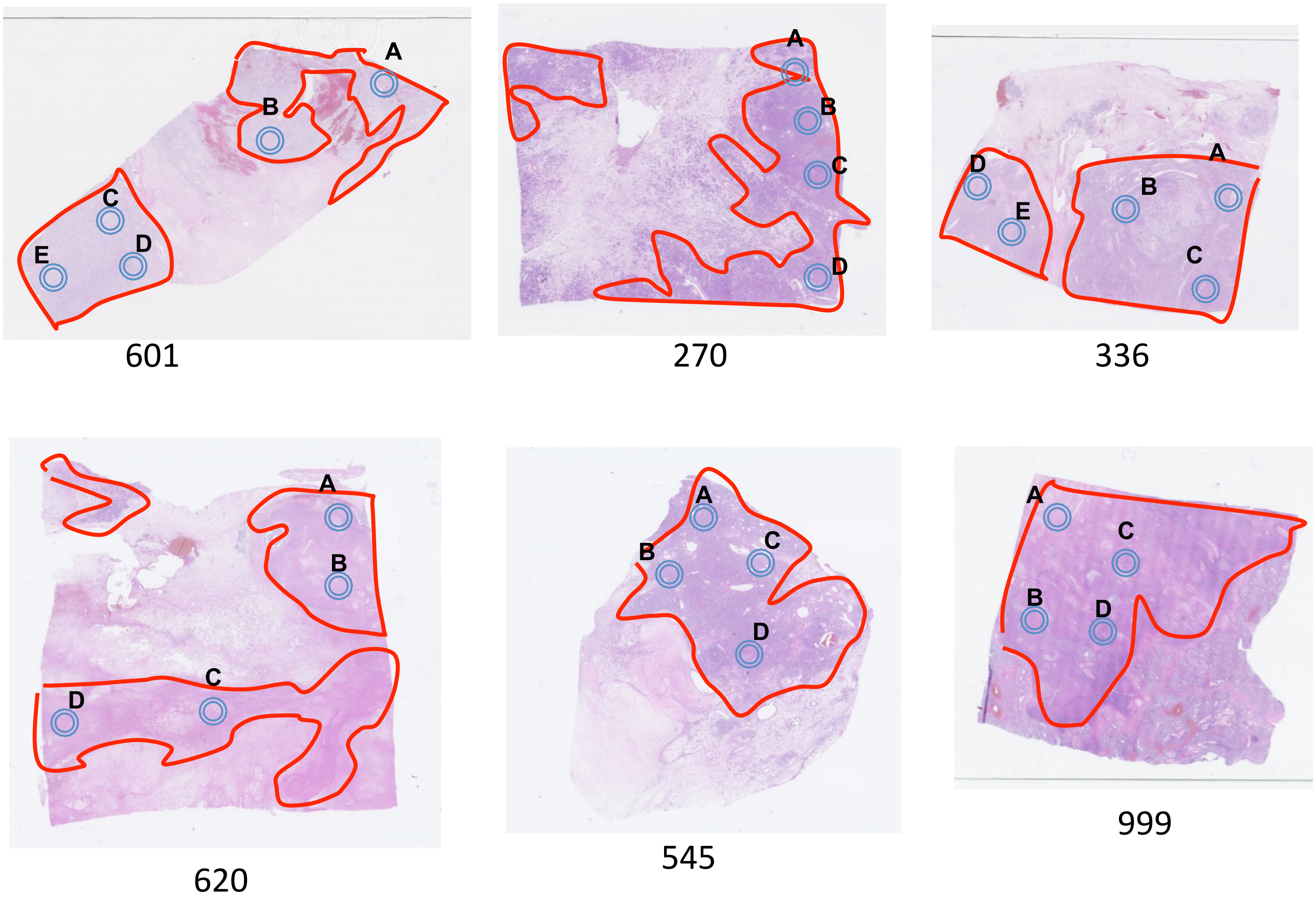

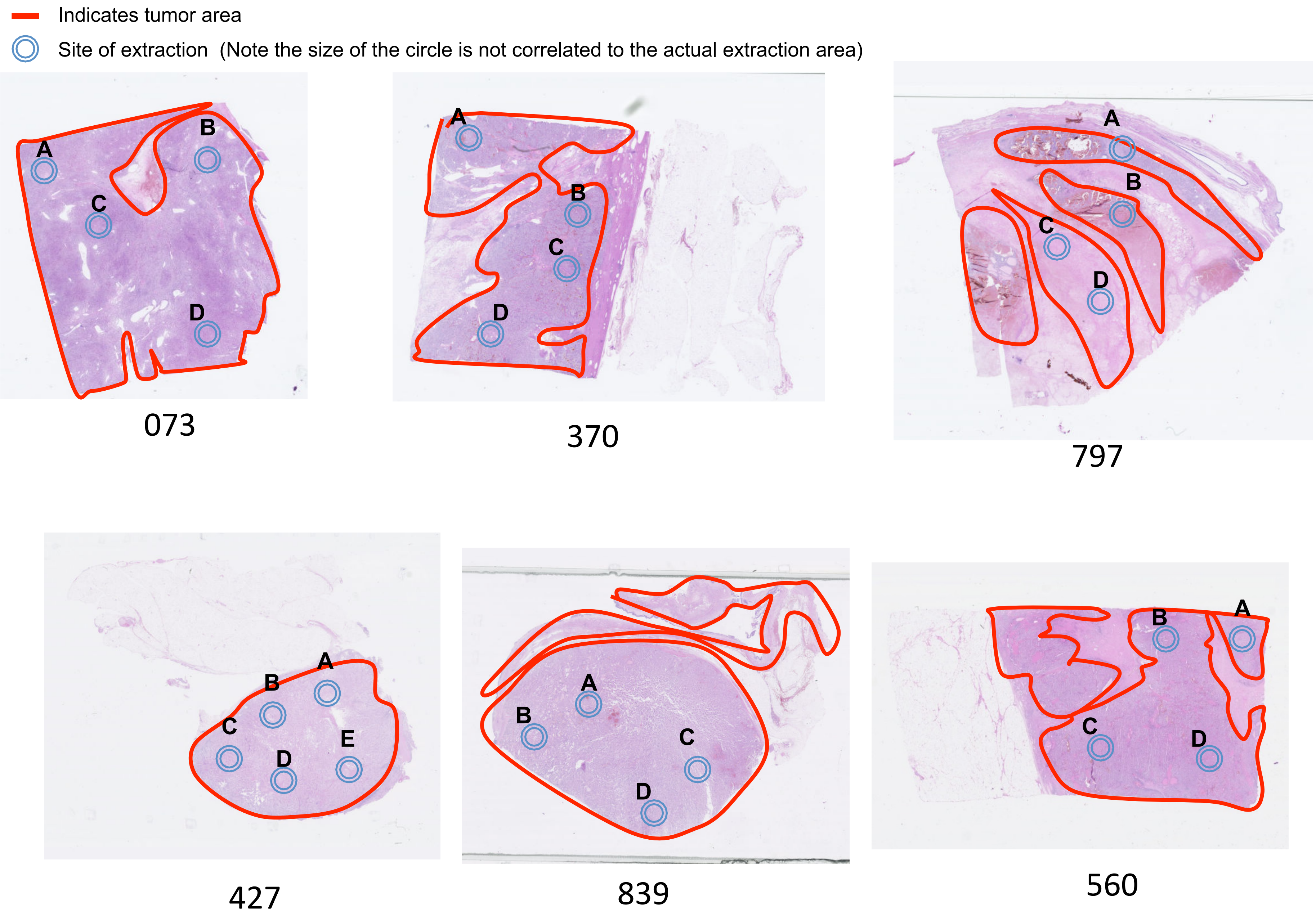
Indicates tumor area
Site of extraction (Note the size of the circle is not correlated to the actual extraction area)
A
B
C
D
073
A
B
C
D
370
A
B
C
D
797
A
B
C
D
E
427
A
B
C
D
839
A
B
C
D
560

Indicates tumor area

Site of extraction (Note the size of the circle is not correlated to the actual extraction area)

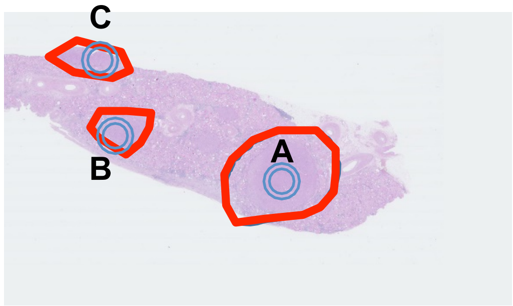


924

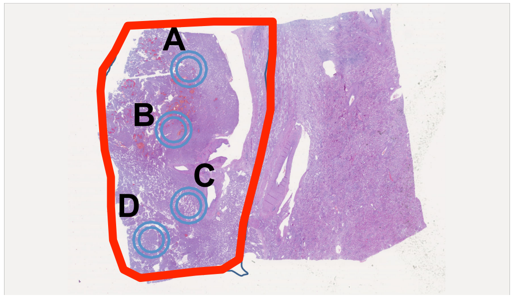


725

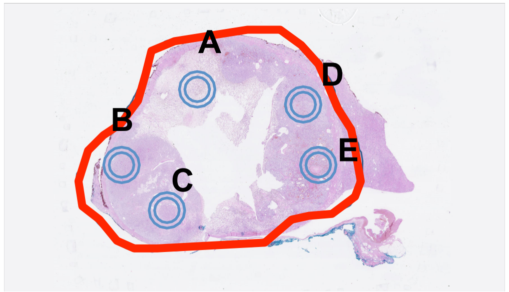


310

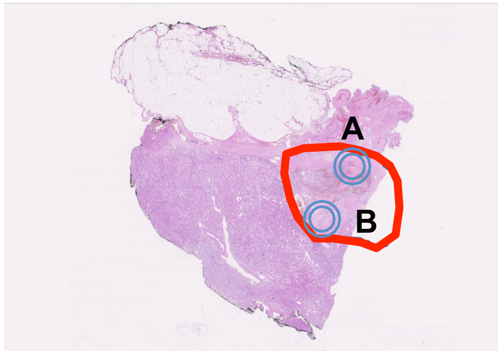


853

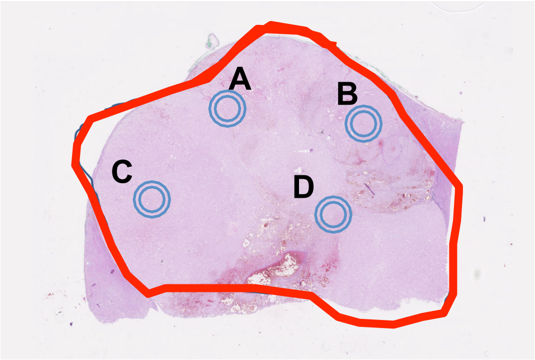


756

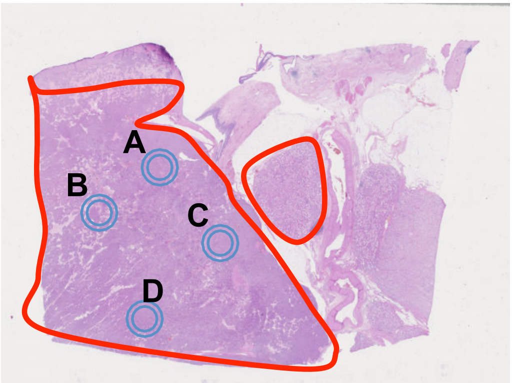


529

Indicates tumor area

Site of extraction (Note the size of the circle is not correlated to the actual extraction area)

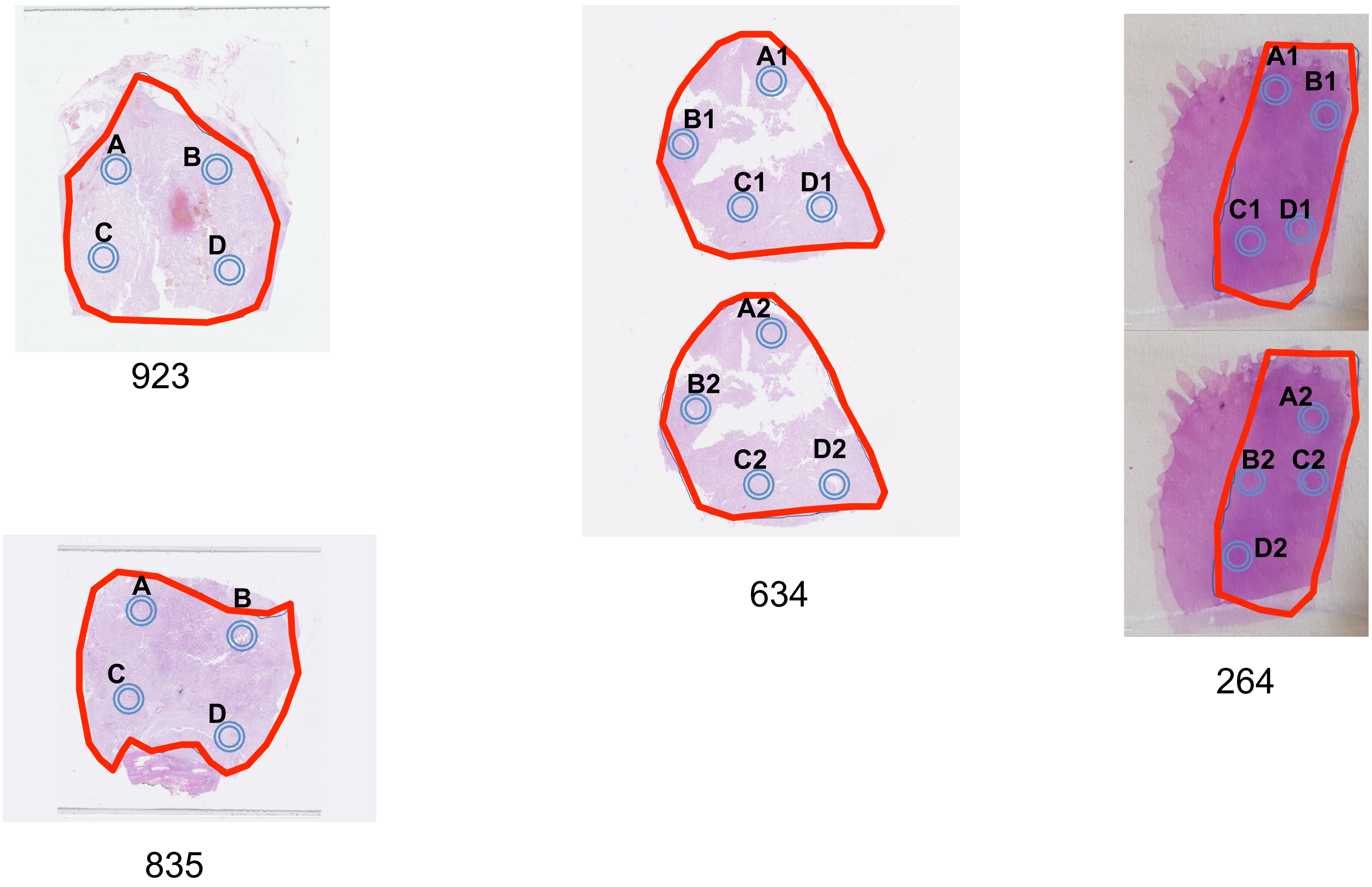

### Supplementary Figure S3: Parameter optimization for PLS-DA

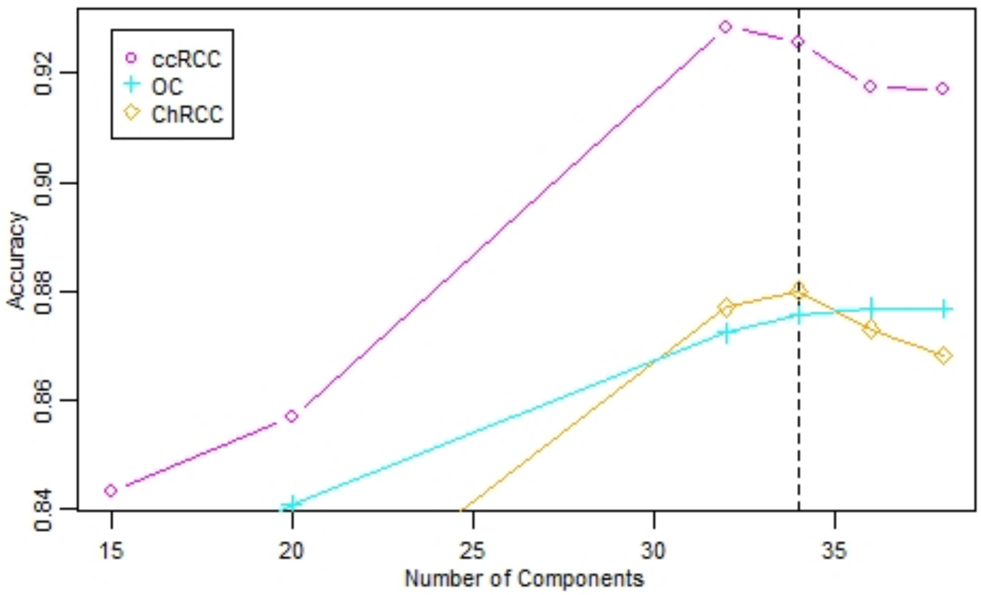


Development of prediction accuracy with different PLS-DA-components. ccRCC RO and ChRCC. Chosen optimum n-components= 34 is marked by dotted line

### Supplementary Figure S4: workflow for imaging data processing

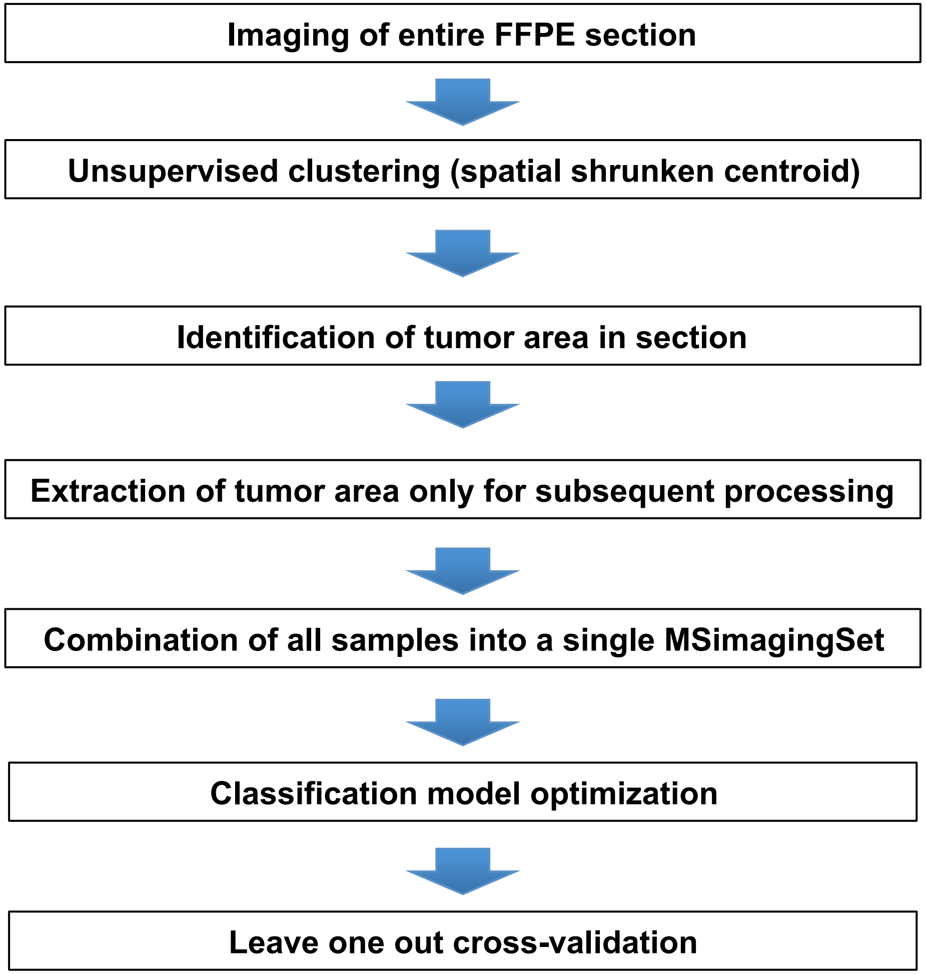


Overview on individual steps of the MALDI-MS imaging classification workflow.

### Supplementary table 3: MSI median fitted scores

| Pannel | Position | Pixel median values: ccRCC | RO | ChRCC | Winner | Pathologist diagnosis | Median absolut deviation: ccRCC | RO | ChRCC | diff to second highest score |
|---|---|---|---|---|---|---|---|---|---|---|
| a | top | 0.1711 | 0.4159 | 0.4113 | RO | RO | 0.1204 | 0.0762 | 0.1474 | 0.0046 |
| | bottom | 0.6520 | 0.0364 | 0.3110 | ccRCC | ccRCC | 0.0627 | 0.1480 | 0.1016 | 0.6156 |
| b | top | 0.0750 | 0.8608 | 0.0657 | RO | RO | 0.1063 | 0.1435 | 0.0560 | 0.7950 |
| | bottom | 0.4987 | 0.4032 | 0.0992 | ccRCC | ccRCC | 0.0789 | 0.0506 | 0.0433 | 0.0954 |
| c | top | -1.0940 | 3.4354 | -1.3545 | RO | RO | 0.9283 | 1.4705 | 0.4987 | 4.7899 |
| | bottom | 2.5791 | -0.3164 | -1.3061 | ccRCC | ccRCC | 0.6037 | 0.6622 | 0.4149 | 2.8955 |
| d | top | -0.0622 | 1.0247 | 0.0378 | RO | RO | 0.1223 | 0.1154 | 0.0403 | 0.9870 |
| | bottom | 1.3893 | -0.3764 | -0.0054 | ccRCC | ccRCC | 0.3803 | 0.3601 | 0.0509 | 1.7658 |
| e | top | 0.3133 | 0.3496 | 0.3363 | RO | RO | 0.0562 | 0.0659 | 0.0716 | 0.0133 |
| | bottom | 0.5013 | 0.3224 | 0.1841 | ccRCC | ccRCC | 0.0588 | 0.0890 | 0.0753 | 0.1789 |
| f | top | 0.3433 | 0.5687 | 0.0865 | RO | RO | 0.1068 | 0.1094 | 0.0231 | 0.4822 |
| | bottom | 1.0971 | 0.0065 | -0.0906 | ccRCC | ccRCC | 0.1471 | 0.0981 | 0.0726 | 1.0906 |
| g | top | -0.6215 | 1.5242 | 0.0451 | RO | RO | 0.4375 | 0.7039 | 0.3552 | 1.4791 |
| | bottom | -0.3673 | 1.6751 | -0.3266 | RO | ccRCC | 0.5359 | 0.6621 | 0.2159 | 2.0423 |
| h | top | 0.0786 | 0.7187 | 0.1868 | RO | RO | 0.1870 | 0.1716 | 0.0616 | 0.5318 |
| | bottom | 0.6651 | -0.0372 | 0.3495 | ccRCC | ccRCC | 0.1936 | 0.1854 | 0.1178 | 0.7023 |
| i | top | 0.2093 | 0.7147 | 0.0811 | RO | RO | 0.1692 | 0.1671 | 0.0388 | 0.6336 |
| | bottom | 0.7213 | 0.2965 | -0.0148 | ccRCC | ccRCC | 0.1304 | 0.1096 | 0.0441 | 0.4248 |
| j | | 0.1030 | 0.2432 | 0.6535 | ChrCC | ChRCC | 0.1058 | 0.1570 | 0.1257 | 0.4104 |
| k | | -0.0056 | 0.4705 | 0.5442 | ChrCC | ChRCC | 0.1396 | 0.1038 | 0.1512 | 0.0737 |
| l | | 0.5634 | -0.3247 | 0.7352 | ChrCC | ChRCC | 0.1649 | 0.2135 | 0.1890 | 0.1718 |
| m | | 0.1427 | 0.2096 | 0.6545 | ChrCC | ChRCC | 0.0709 | 0.1466 | 0.1636 | 0.4450 |
| n | | 0.2312 | 0.3089 | 0.4746 | ChrCC | ChRCC | 0.1461 | 0.1328 | 0.1374 | 0.1656 |

misassignment
differnce<0.05

Median values of classification fitted scores. Panel column refers to panels in Figure 3.
Top and bottom indicate sample positions within the panels of figure 3.

#### Supplementary Figure S5: boxplot representation of scores from PLS-DA classification

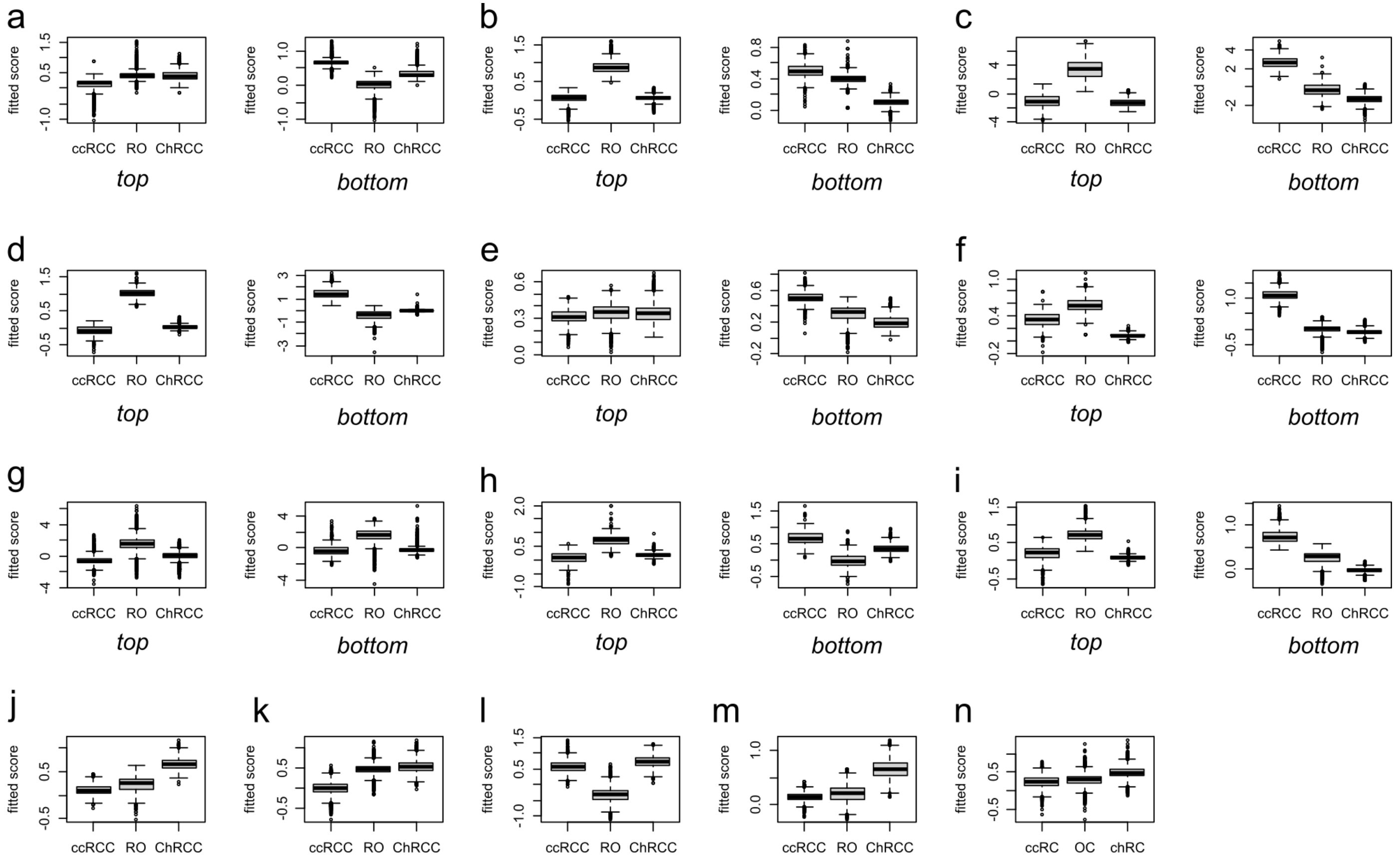


Boxplot of PLS-DA fitted scores across all pixels in a patient sample. Scores for each individual testing condition (ccRCC, RO, ChRCC) for a given patient sample are plotted in one panel. S5 a-n) correlate with panels 3 a-n) in Figure 3

#### **Supplementary Figure S6: PLS coefficients as a function of *m/z*.**

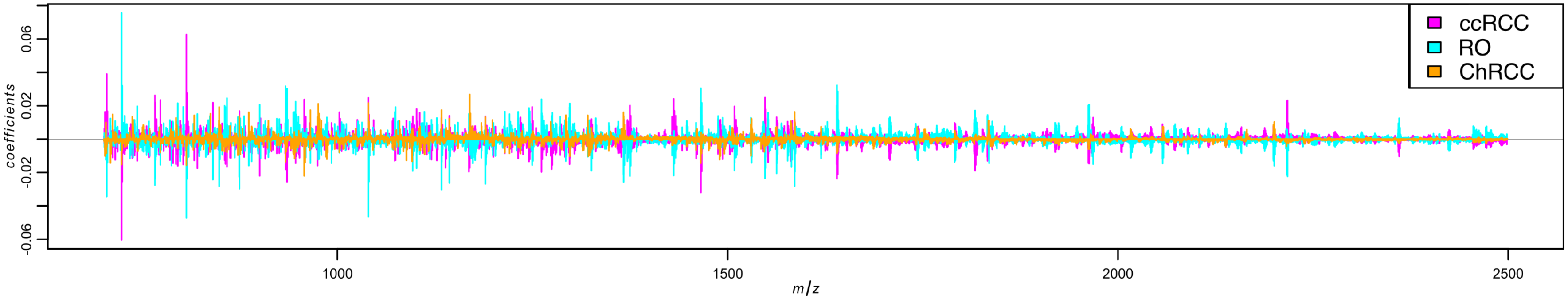


The diagram displays the impact of each detected *m/z* signal feature in imaging MS data from PLS-DA prediction (spectra are binned to 0.25 m/z bins). Positive coefficient indicates presence or higher abundance in the respective condition. Negative coefficient indicates absence or lower abundance of the m/z value in the respective condition (a list with the 100 most influential features can be found in supplementary material 2)

### Supplementary Figure S7: Unprocessed version of Figure 3

Cross-validation results of PLS-DA classification

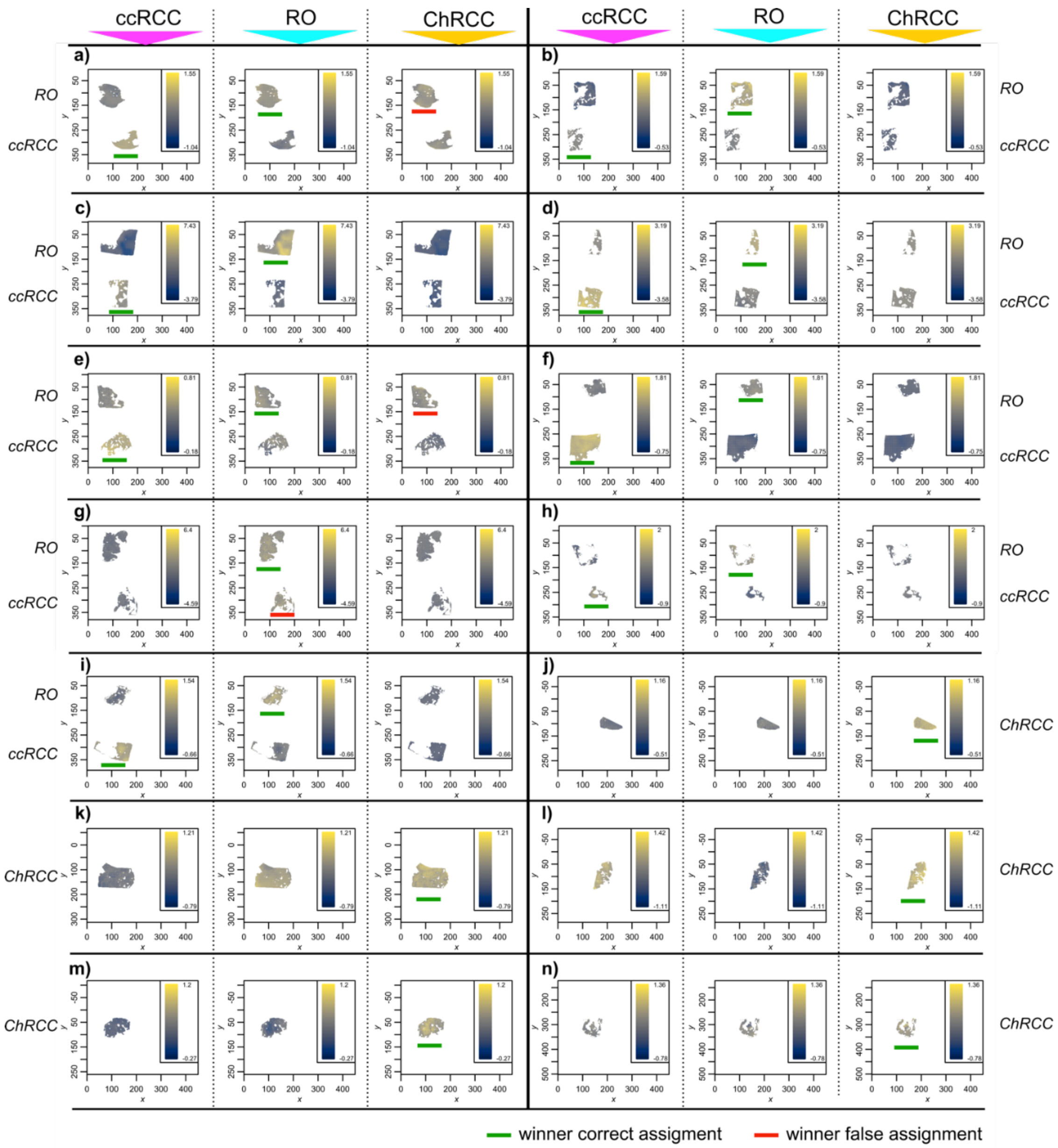
